# Mycorrhizal fungi modulate tree diversity effects on nutrient dynamics

**DOI:** 10.1101/2023.12.06.569218

**Authors:** Elisabeth Bönisch, Evgenia Blagodatskaya, Rodolfo Dirzo, Olga Ferlian, Andreas Fichtner, Yuanyuan Huang, Samuel J. Leonard, Fernando T. Maestre, Goddert von Oheimb, Tama Ray, Nico Eisenhauer

**Affiliations:** German Centre for Integrative Biodiversity Research (iDiv) Halle-Jena-Leipzig, Puschstr. 4, 04103 Leipzig, Germany; Institute of Biology, Leipzig University, Leipzig, Germany; Helmholtz-Centre for Environmental Research (UFZ) Soil Ecology Department, Theodor-Lieser-Str. 11, 06120 Halle, Germany UFZ; Departments of Biology and Earth Systems Science, Stanford University, Stanford, CA, USA 94305; Institute of Ecology, Leuphana University of Lüneburg, Universitätsallee 1, 21335 Lüneburg, Germany; Instituto Multidisciplinar para el Estudio del Medio “Ramón Margalef”, Universidad de Alicante, Carretera de San Vicente del Raspeig s/n, Alicante, Spain; Departamento de Ecología, Universidad de Alicante, Carretera de San Vicente del Raspeig s/n, Alicante, Spain; Institute of General Ecology and Environmental Protection, TU Dresden University of Technology, Pienner Straße 7, 01737 Tharandt, Germany; Institute of Biology/Geobotany and Botanical Garden, Martin Luther University Halle-Wittenberg, 06108 Halle (Saale), Germany

**Keywords:** *diversity effects*, *mycorrhizal fungi*, *MyDiv*, *nutrient dynamics*, *plant-soil interaction*, *tree species richness*

## Abstract

Species-specific differences in nutrient acquisition strategies allow for complementary use of resources among plants in mixtures, which may be further shaped by mycorrhizal associations. However, empirical evidence of these relationships is scarce, particularly for tree communities.

We investigated the impact of tree species richness and mycorrhizal types, arbuscular mycorrhizal fungi (AM) and ectomycorrhizal fungi (EM), on the above- and belowground carbon (C), nitrogen (N), and phosphorus (P) dynamics.

Soil and microbial biomass elemental pools did not strongly respond to tree species richness or mycorrhizal type. Tree species richness increased foliage C and P pools depending on mycorrhizal type. Additive partitioning analyses showed that net biodiversity effects for C, N, P pools in EM tree communities, and N pools in AM tree communities, were driven by selection effects, while mixtures of both mycorrhizal types were influenced by complementarity effects. Furthermore, tree species richness enhanced soil nitrate uptake over two years but had no impact on ammonium and phosphate levels.

Our results indicate that positive effects of tree diversity on aboveground nutrient storage are mediated by complementary mycorrhizal strategies. Given the prevalence of anthropogenic impacts on tree species richness globally, these results may have important implications for reforestation of multifunctional forests.

## Introduction

The positive relationship between biodiversity and ecosystem functions in terrestrial ecosystems (e.g. Cardinale *et al*., 2007; Morin *et al*., 2011; Huang *et al*., 2018) largely depends on plant nutrient availability, uptake, and their spatio-temporal dynamics (Barry *et al*., 2019). Theory predicts that these effects are substantially driven by dissimilarities in resource acquisition traits (i.e. different rooting systems; (Dornbush & Wilsey, 2010) and resource use strategies of plant species (i.e. conservative vs. acquisitive; Diaz *et al*., 2004; Barry *et al*., 2019). Diverse species assemblages occupy a greater number of resource niches which reduces competition for limiting nutrients (Loreau & Hector, 2001; Hooper *et al*., 2005; Ferlian *et al*., 2018). The complementary use of resources is expected to enhance resource use and net primary productivity of a community (Tilman, 1980).

Besides physiological differences between species supporting resource use complementarity (Barry *et al*., 2019), plants associate with important symbiotic partners – mycorrhizal fungi (van der Heijden *et al*., 1998). Mycorrhizal fungi support their plant hosts directly by supplying them with water, mineral nutrients, and protecting them against pathogens (Smith & Read, 2008), as well as indirectly by improving biogeochemical soil properties, such as soil structure and pH (Rillig & Mummey, 2006). In return, plants provide the fungi with carbohydrates and lipids (Luginbuehl *et al*., 2017). Trees predominantly associate with either arbuscular mycorrhizal fungi (AM) or ectomycorrhizal fungi (EM) or both, and these two mycorrhizal types can have different effects on resource uptake processes (Bonfante & Genre, 2010; Ferlian *et al*., 2021). AM fungi strongly support the provision of phosphorus to plants (Smith & Smith, 2011) via resource acquisitive traits, characterized by fast biochemical cycling, rapid growth of the host plant (Powell & Rillig, 2018), and nutrient-rich soils, foliage and litter (Phillips *et al*., 2013; Deng *et al*., 2023). EM fungi enhance the availability and uptake of organic nutrients, mainly organic nitrogen (van der Heijden *et al*., 1998) to plants by directly promoting the decomposition of organic matter through the exudation of extracellular enzymes (Diaz *et al*., 2004; Lambers *et al*., 2009; Phillips *et al*., 2013). However, this favors a competitive condition between EM and free-living decomposers in soil, slowing down decomposition of plant litter (Averill & Hawkes, 2016) thereby promoting carbon accumulation in soils (Orwin *et al*., 2011; Averill & Hawkes, 2016). This slow cycling system (conservative resource use strategy) is characterized by slow growth and slow foraging roots of host plants, poor quality foliage and litter (Cornelissen *et al*., 2001; Midgley & Phillips, 2014; Averill & Hawkes, 2016; Eskelinen *et al*., 2020). Thus, the presence of two distinct mycorrhizal types, characterized by dissimilar resource uptake strategies and traits, is expected to enhance resource complementarity (Cheng *et al*., 2016) among their associated plant hosts and alter above- and belowground elemental contents and pools (Ferlian *et al*., 2018; Eisenhauer *et al*., 2022).

Mycorrhizal fungi, as well as plant diversity, affect overall soil microbial communities. Studies confirmed the positive effect of tree species richness on diversity and composition of soil microbes (Bardgett & van der Putten, 2014; Deng *et al*., 2023), signaling that resource complementarity between different tree species plays a crucial part (Singavarapu *et al*., 2021). Due to their distinct functional roles, mycorrhizal fungi strongly affect other soil microbes (Powell & Rillig, 2018). For example, high-quality litter (low carbon-to-nitrogen ratio) of AM trees generally enhances microbial activity and release of macronutrients in soil (Aislabie & Deslippe, 2013; Bardgett & van der Putten, 2014), while the low-quality litter of EM trees triggers competition for nutrients between the fungi and other microbial decomposers (Averill *et al*., 2014; Averill & Hawkes, 2016).

While resource complementarity is expected to play a significant role in space, via the occupation of different nutrient niches in soil by roots, mycorrhizal hyphae, and other microbial communities, nutrient availability is also highly dynamic in time (Eisenhauer *et al*., 2022). Availability of nutrients is strongly affected by seasonal changes of environmental factors, such as temperature, precipitation, and soil moisture (Niklaus *et al*., 2001; Scherer-Lorenzen *et al*., 2003; Oelmann *et al*., 2007). This is caused by the dependency of soil microorganism activity on suitable climatic conditions (Kuzyakov & Blagodatskaya, 2015; Baldrian, 2017). Therefore, mycorrhizal fungi also exhibit a distinct seasonality (Keller & Phillips, 2018), which may further affect their ability to supply their plant hosts with nutrients. Moreover, both mycorrhizal strategies may complement each other over time and therefore enhance total nutrient uptake. However, direct effects of mycorrhizal types and tree species richness on the seasonal availability of nutrients have not been studied yet.

Here we aim to shed light on the patterns of nutrient dynamics mediated by mycorrhizal type and tree species richness. The research was conducted within the tree diversity experiment MyDiv, which manipulates both tree species richness and mycorrhizal strategy (Ferlian *et al*., 2018). Knowledge of spatial and temporal nutrient dynamics may improve our understanding of the underlying mechanism driving positive diversity effects. This motivated us to test the following hypotheses:

### H 1

Both species-rich and tree communities with mixed mycorrhizal types (AM+EM) will have higher carbon (C), nitrogen (N), and phosphorus (P) contents in soil, soil microbial biomass, and foliage. Consequently, these tree communities will have the largest elemental pools.

### H 2

Communities with increased tree species richness and mixed mycorrhizal strategies will enhance resource exploitation from soils and therefore show lower contents of plant-available soil nutrients. These effects will be more pronounced during the growing season and under favorable microclimatic conditions, such as increased soil moisture.

### H 3

We expect positive effects of high tree species richness and mixed mycorrhizal types (AM+EM) on aboveground elemental pools to be explained by higher complementarity between tree species and mycorrhizal types.

To test these hypotheses, we characterized the predominantly dynamic above- and belowground chemical properties (soil, soil microbial biomass, and foliage carbon (C), nitrogen (N), and phosphorus (P) contents and respective pools) and temporal availability of the nutrients nitrate (NO_3_-N), ammonium (NH_4_-N), and phosphate (PO_4_-P) in the soil for two years.

## Materials and Methods

### Study site and experimental design

MyDiv is a long-term tree diversity experiment, located in Bad Lauchstädt (Saxony-Anhalt, Germany). This experiment is run by the German Center of Biodiversity Research (iDiv), Halle-Jena-Leipzig and Helmholtz Center for Environmental Research (UFZ) is part of TreeDivNet (www.treedivnet.ugent.be). The experiment was established in March 2015 on former arable land having a nutrient-and humus-rich Chernozem soil that is rich in N, but limited in P (Ferlian *et al*., 2018). The experimental site is divided into two blocks according to a gradient in abiotic parameters determined before the establishment (Ferlian *et al*., 2018). The experiment consists of 80 plots in total. Each of the 11 m x 11 m plots is divided into a buffer zone and a core zone (64 m²) and covered by a weed tarp. Per plot, 140 two-year old tree saplings were planted with 1 m distance from each other (64 trees in the core zone). The species pool contains 10 deciduous tree species, five of which associate predominantly with AM and five with EM fungi (Table 1). The selected deciduous tree species are native to Germany, adapted to the site conditions, and are of either economical or recreational relevance (Ferlian *et al*., 2018). Further, only one species per genus was selected to have representative species widely spread across the angiosperm phylogeny (Ferlian *et al*., 2018). However, based on these selection criteria, we could not avoid that four of the five EM species belong to the order Fagales. The experiment combines three levels of tree species richness (monocultures, two-species mixtures, and four-species mixtures) with three levels of mycorrhization, either AM, EM, or a mixture of both (plot-specific details on experimental design can be found in Supporting Information Table S1). The treatments of the experiment were confirmed by DNA-sequencing and determining mycorrhization rates, indeed demonstrating that AM trees had substantially higher mycorrhization rates by arbuscular mycorrhizal fungi than by ectomycorrhizal fungi, whereas the opposite pattern was found for EM trees (Table S1 in Ferlian *et al*. 2021). Furthermore, AM and EM richness significantly increased with tree species richness (Ferlian *et al*., 2021).

**Table 1.** Overview of tree species used in the MyDiv experiment with their respective mycorrhizal association (arbuscular mycorrhizal fungi, AM; ectomycorrhizal fungi, EM). For details, see Ferlian *et al*. (2018).

| Species | Family | Mycorrhizal association |
| --- | --- | --- |
| *Acer pseudoplatanus* L. | Sapindaceae | AM |
| *Aesculus hippocastanum* L. | Sapindaceae | AM |
| *Fraxinus excelsior* L. | Oleaceae | AM |
| *Prunus avium* L. | Rosaceae | AM |
| *Sorbus aucuparia* L. | Rosaceae | AM |
| *Betula pendula* Roth | Betulaceae | EM |
| *Carpinus betulus* L. | Betulaceae | EM |
| *Fagus sylvatica* L. | Fagaceae | EM |
| *Quercus petraea* (Matt.) Liebl. | Fagaceae | EM |
| *Tilia platyphyllos* Scop. | Malvaceae | EM |

### Data collection

#### Soil physico-chemical measurements

Soil samples for the measurement of soil bulk density (g cm^-3^) were collected with a 5 cm diameter soil corer to a depth of 5 cm in May 2021. The samples were air dried, weighted, and the core volume calculated. The bulk density was then calculated as the dry weight of soil divided by its volume. Soil samples for the analyses of soil pH were taken with 5 cm soil corers to 10 cm soil depth in October 2021. Four cores were taken per plot, where all tree species were equally represented as neighbors, and finally pooled to one composite sample per plot. Plant material and stones were removed by sieving (2 mm mesh). For pH measurements, soil samples were solved in 25 ml CaCl_2_-solution (0.01 mol l^-1^), stirred, left for one hour, stirred again, and measured with the probe.

For the assessment of soil C, N, and P contents (g kg^-1^), samples were taken with 5 cm soil corers to 10 cm depth in October 2021. This approach is in line with common soil biodiversity and function monitoring approaches (e.g. Guerra *et al*., 2021a,b) and based on the biological activity of plants and animals which is highest in the first 5-10 centimeters of the soil and decreases with deeper layers (e.g. Stone *et al*., 2014; Weldmichael *et al*., 2020). Four cores were taken per plot, where all tree species were equally represented as neighbors, and finally pooled to one composite sample per plot. Plant material and stones were removed by sieving (2 mm mesh). For analysis of total C, N, and P, samples were dried at 60 °C, ground to fine powder with a ball mill (MM 400, Retsch, Haan, Germany). For C and N analysis, samples were dried for another 24 h, and transferred into tin capsules. Analyses were conducted using an elemental analyzer (VarioMax, Elementar Analysensysteme GmbH, Langenselbold). The analysis of P was conducted by dissolving an aliquot of 500 mg ground soil using microwave digestion at 200°C for 30 minutes (5 ml HNO_3_ and 0.5 ml H_2_O_2_ to avoid production of nitric oxides; Multiwave, Anton Paar GmbH, Graz, Austria). Measurements were carried out with an inductively coupled plasma optical emission spectrometer (wavelength: 177.5 nm; limit of determination: 0.13 mg l^-1^; Arcos, Spectro Analytical Instruments GmbH, Kleve, Germany).

#### Soil microbial C, N, and P measurements

For the analysis of microbial C, N, P contents (μg g^-1^), soil samples (same pooled samples as for soil C, N, and P analyses) were kept at 4°C and analyzed within three days after sampling by chloroform-fumigation-extraction method (CFE) using 0.05 M K_2_SO_4_ extracts and a conversion factor (Kp) of 0.45 for C and 0.54 for N (Brookes *et al*., 1985; Wu *et al*., 1990). Microbial P content was determined by direct fumigation and anion exchange membrane techniques (Yevdokimov *et al*., 2016) with conversion factor 0.40 (Brookes *et al*., 1985). We also tested the calculation of a conversion factor according to the soil properties (Bilyera *et al*., 2018), and confirmed that the Kp values for P did not differ significantly from 0.40 for the soils used in our study.

#### Time series of available soil N and P

To assess the intra-annual and inter-annual variation, as well as seasonal differences in soil nitrate (NO_3_-N), ammonium (NH_4_-N), and phosphate (PO_4_-P), we inserted ionic exchange membranes (IEM) each month. IEMs are a valid replacement of traditional methods to measure in situ nutrient availability (measured as μg cm-2 day-1) for plants in soil. Experimental studies show that ion contents obtained by IEMs correlate with plant uptake of such ions (Ziadi *et al*., 1999; Qian & Schoenau, 2002; Durán *et al*., 2013). For two years (April 2019 to March 2021), cationic and anionic IEMs were inserted into the soil at 10 cm depth and replicated five times in a transect of 1 m per plot each month. After incubation for one month, the IEMs were dried at room temperature and cleaned. Further processing of the samples was conducted according to (Rodríguez *et al*., 2009; Durán *et al*., 2013). To assess daily contents of plant-available nutrients, the measured nutrient contents were divided by the days of incubation.

#### Foliage C, N, and P measurement

For the analysis of foliar C, N, and P contents (g kg^-1^), fresh leaves of all tree species were sampled in July 2021. Nine leaves from three different tree individuals per plot and tree species were sampled at a tree height of approximately 2.5 m. The leaves were dried at 40 °C in the drying oven for 24 h and ground with a ball mill (MM 400, Retsch, Haan, Germany). For C and N analyses, samples were transferred into tin capsules and analyzed using an elemental analyzer (Vario EL cube; Elementar Analysensysteme GmbH, Langenselbold). The analysis of foliage P was conducted the same way as for the soil samples.

#### Environmental data

Environmental data (soil moisture (%) and soil temperature (°C) in 10 cm depth; air temperature (°C), air humidity (%)), were measured with three weather stations (mean value used) at the MyDiv experimental site (Meteorological data of Bad Lauchstädt, Helmholtz Centre for Environmental Research (UFZ), Department of Soil System Science). Therefore, only site-specific data were available in 2021, but not plot specific data.

### Calculations

#### Soil, soil microbial, and foliage pools

For each plot, soil nutrient pools (g m^-2^) and soil microbial biomass nutrient pools (g m^-2^) were calculated as the products of total C, N, and P and soil bulk density (g cm^-3^) (Rochow, 1975). Community foliage elemental pools (per plot) were derived by multiplying foliage elemental contents (g kg^-1^) with an index that captures the spatial structural complexity of biomass distribution within a stand (Stand Structural Complexity Index, SSCI; (Ehbrecht *et al*., 2017), since data on individual leaf biomass or Leaf Area Index (LAI) were not available. The SSCI is based on terrestrial laser scanning data, which were collected in September 2021 at the study site (see Ray *et al*., 2023). Here, we use the SSCI as a proxy for LAI assuming that structurally more complex stands are associated with greater spatial complementarity in canopy space, and thus greater light interception (Ray *et al*., 2023). For example, differences in branching intensity and branch density lead to greater crown complementarity (Hildebrand *et al*., 2021), which in turn should result in a higher leaf foliage production of structurally more complex stands.

#### Biodiversity effects

Net biodiversity effects, selection effects, and complementarity effects for the elemental pools were calculated using the additive partitioning method of Loreau & Hector (2001), based on the following equations.

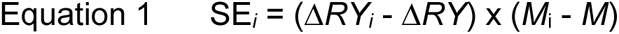

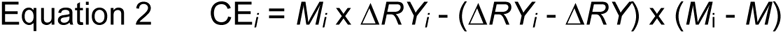

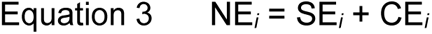

Here, Δ*RY_i_* represents the deviation from the expected relative yield of species *i* in the two- and four-species mixture (*RY*_observed_-*RY*_expected_), and Δ*RY* represents the average relative yield deviation of all species in the tree stand. *M_i_* represents the yield of species *i* in the monocultures, and *M* is explained as the average yield of all species in the monoculture. For this, the species-specific elemental pools (Supporting Information Fig. S3) were estimated from the community pools by using species-specific wood volume (m^3^ m^-2^) (Eqn. 4; Supporting Information Fig. S1, S2; Table S2, S3).

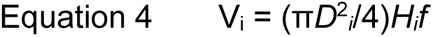

where *D_i_* represents the stem diameter (m) for each tree *i* measured 5 cm above ground and *H_i_* is the tree height (m). To account for the deviation of the tree volume from the volume of a cylinder *f* is added as a cylindrical form factor of 0.5 (Ray *et al*., 2023). Dead trees (2.6 % according to the tree inventory data from 2021) were not considered in the analysis.

### Statistical Analyses

Linear mixed-effects models (Type I Sum of Squares) were used to assess the impact of tree species richness (numerical with three levels - one, two, four), mycorrhizal type (factorial, three levels: AM, EM, AM+EM), and their interacting effect on elemental contents, elemental pools, and biodiversity effects (net biodiversity effects, selection effects, complementarity effects). Block (factorial, two levels: one, two) was used as a random effect. Further, effects of tree species richness and mycorrhizal type on soil pH and soil bulk density were tested (results can be found in Supporting Information Table S4). To test for effects between groups (e.g. AM vs. EM; AM+EM vs. AM; AM+EM vs. EM) Tukey’s honestly significant difference (HSD) tests were conducted. The relationship between response variables (C, N, P contents and C, N, P pools of soil, soil microbial biomass, and foliage; biodiversity effects) and tree species richness as single explanatory variable for each group (AM, EM, AM+EM) was tested with simple linear regression analysis. We used the percentage change as a standardized measure to quantify the strength of biodiversity effects (tree species richness, mycorrhizal type) on the response variables (C, N, P contents and pools) (Supporting Information Table S15 – S18).

To improve the normality of residuals, biodiversity effects were square-root-transformed with sign reconstruction (sign(y) = |y|) (Loreau & Hector, 2001). To test whether biodiversity effects were larger than zero, we used one-tailed *t*-tests. Significantly larger biodiversity effects indicate overperformance in two- and four-species tree communities relative to monocultures.

For the analysis of the time series dataset on plant available nutrients Bayesian statistics was used to include the temporal trend to estimate the parameters of the models. Models were fitted using R-INLA (R-Integrated Nested Laplace Approximation), with tree species richness, mycorrhizal type, season and soil moisture, as well as their interactions as fixed effects, and block, plot and year as random effects. To account for the temporal trend, we added a RW1 (Random walk model of order 1) trend. For model comparison DIC values were used and insignificant interaction terms removed. The final model is given below.

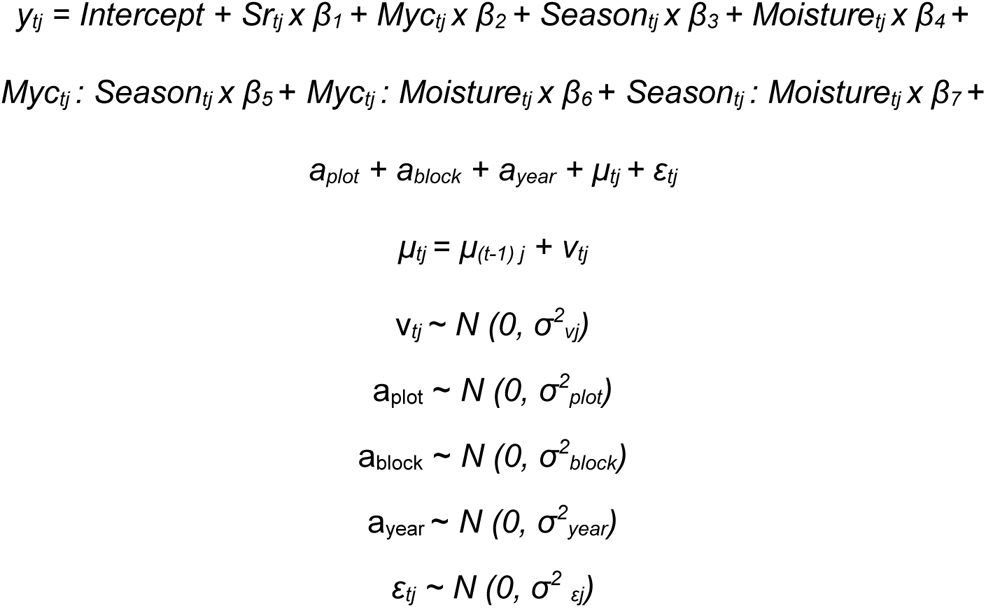

Here, *y_tj_* represents the response variable soil nitrate (NO_3_-N), ammonium (NH_4_-N), or phosphate (PO_4_-P) at time *t* for plot *j*. *Sr_tj_, Myc_tj_, Season_tj_,* and *Moisture_tj_* indicate tree species richness, mycorrhizal type, season and soil moisture at time t for plot j respectively, while *a_plot_, a_block_,* and *a_year_* stand for the random effects of plot, block and year respectively. We assume that each random effect is independent and identical distributed with a mean of zero and its variance. The trend *μ_tj_* is modelled as a RW1 random walk trend based on penalized complexity (Zuur *et al*., 2017) prior with parameters of U = 1 and *α* = 0.01. Here, we allowed separate trends for each tree species richness and each mycorrhizal type combination within each year.

All statistical analyses were performed using R Statistical Software (version 4.3; R Development Core Team, http://www.R-project.org) using the packages *lme4* (version 1.1-33; (Bates *et al*., 2015) and *emmeans* (version 1.8.6; (Lenth *et al*., 2023) for mixed-effects model analysis and Tukey’s HSD tests, respectively. For the additive partitioning method (calculation of biodiversity effects) the function *addpart* from the package *pdiv* was used (Niklaus 2022). For the Bayesian-based time series analysis *R-INLA* (version 22.12.16) was used (Zuur *et al*., 2017).

## Results

### Elemental contents of foliage, bulk soil, and soil microbial biomass

Overall, effects of tree species richness and mycorrhizal type were most pronounced for the elemental contents of tree foliage. In contrast, no effects of either treatments were observed in bulk soil (Fig. 1d-f; Supporting Information Table S5 b), and they were rarely present in soil microbial biomass (Fig. 1g-i; Supporting Information Table S5 c). Soil microbial biomass N (*p* = 0.039) and P contents (*p* = 0.018) were significantly affected by tree species richness and increased by around 3% and 15% in four-species tree stands compared to monocultures respectively, while C content (*p* = 0.093) was marginally affected. Mycorrhizal strategies did not affect microbial biomass.

**Fig. 1.**
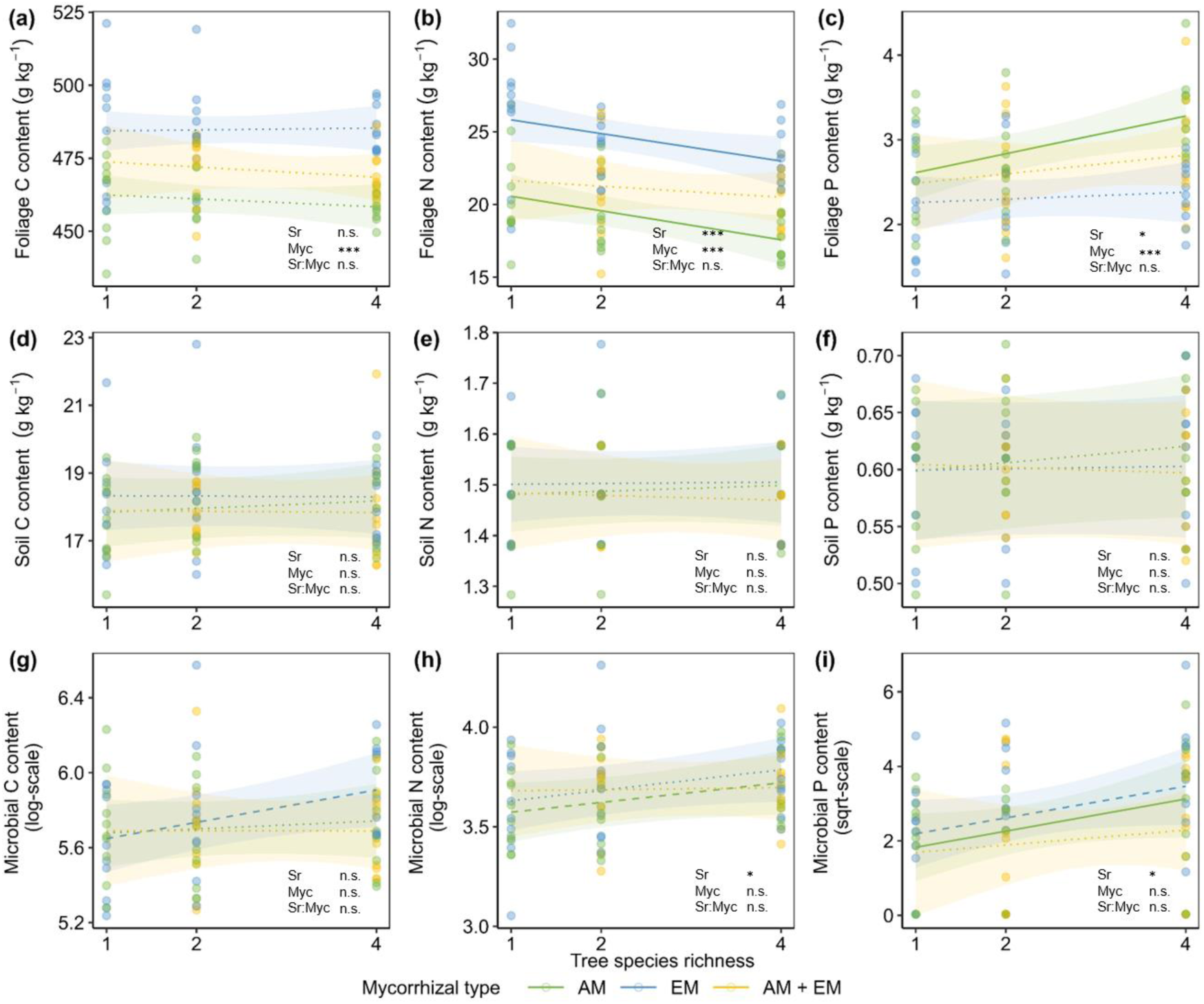
Carbon (C), nitrogen (N) and phosphorus (P) contents in (a-c) foliage, (d-f) soil and (g-i) microbial biomass as a function of Tree species richness (one, two, four; Sr) for communities containing arbuscular mycorrhizal tree species (AM), ectomycorrhizal tree species (EM), or both (AM + EM) tree species (Myc). Each dot represents a tree community, and colors indicate different Mycorrhizal types. Regression lines are based on mixed-effect models (predicted means). Solid lines indicate statistically significant relationships (*p* < 0.05), dashed lines marginally significant relationships (0.1 > *p* > 0.05), and dotted lines non-significant relationships (*p* > 0.1). Statistical significance of main effects is indicated in each panel (n.s., *p* > 0.05; * *p* < 0.05; ** *p* <0.01; *** *p* <0.001).

In tree foliage, tree species richness had contrasting effects on contents of N and P: while N contents decreased significantly with tree species richness (*p* <.001) by about 13% in monocultures compared to four-species communities, P contents increased significantly (*p* = 0.017) by 14% in four-species stands compared to monocultures (Fig. 1a-c; Supporting Information Table S5 a). C contents remained constant along the tree diversity gradient. Mycorrhizal type significantly affected C, N, and P contents (*p* <.001 respectively), whereby C and N contents were highest for EM tree stands (5% and 24% larger compared to AM tree stands, respectively), and P contents for AM tree communities (23% higher compared to EM tree stands; Supporting Information Table S7, S9). Tree stands with mixed mycorrhizal types showed additive effects. We found no interaction effect of mycorrhizal type and tree species richness on elemental contents.

### Elemental pools of foliage, bulk soil, and soil microbial biomass

We found only effects of tree species richness and mycorrhizal type on elemental pools of tree foliage (Fig. 2, Supporting Information Table S6), but not on pools of bulk soil and soil microbial biomass (Supporting Information Fig. S4; Table S6). Tree species richness had an overall positive effect on C (*p* <.001) and P (*p* <.001) pools in foliage, whereby the pool sizes increased by about 25% and 41%, respectively, in four-species mixtures compared to monocultures (Supporting Information Table S10, S16). While N pools in foliage were not significantly affected by tree species richness, they were significantly affected by mycorrhizal type (p <.001), whereby N pools in EM tree stands were about 28% larger compared to N pools in AM tree communities. In contrast, AM tree communities obtained 18% larger P pools in foliage compared to EM tree stands (Supporting Information Table S8). However, no significant main effect of mycorrhizal type on P and C pools was found (Supporting Information Table S6). In addition, the difference between pool sizes of EM and AM tended to increase as species richness increased. For mixtures of the two mycorrhizal types, we consistently observed additive effects, similar to the elemental contents. Significant interaction effects between the two treatments were not detected.

**Fig. 2.**
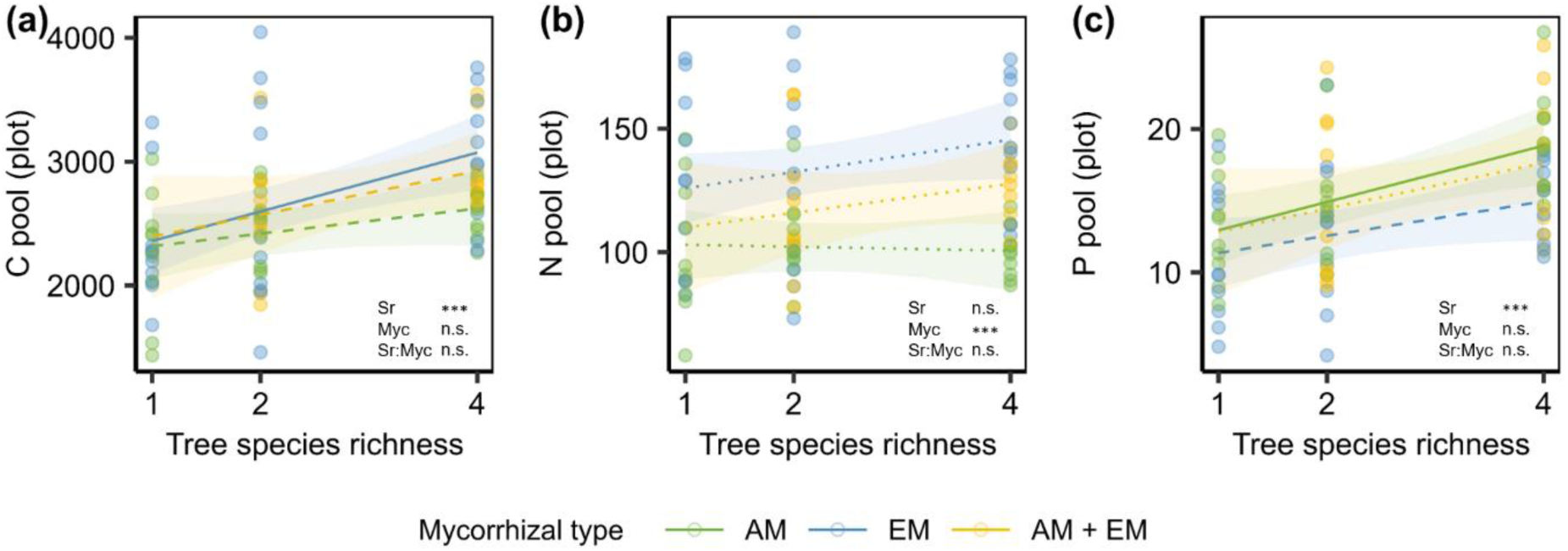
Foliage (a) carbon (C), (b) nitrogen (N), and (c) phosphorus (P) pools as affected by Tree species richness (one, two, four; Sr) for communities containing arbuscular mycorrhizal tree species (AM), ectomycorrhizal tree species (EM), or both (AM + EM) tree species (Myc). Each dot represents a tree community, and colors indicate different Mycorrhizal types. Regression lines are based on mixed-effect models (predicted means). Solid lines indicate statistically significant relationships (*p* < 0.05), dashed lines marginally significant relationships (0.1 > *p* > 0.05), and dotted lines non-significant relationships (*p* > 0.1). Statistical significance of main effects is indicated in each panel (n.s., *p* > 0.05; * *p* < 0.05; ** *p* <0.01; *** *p* <0.001).

### Seasonal availability of soil nitrate, ammonium, and phosphate

Our study reveals pronounced temporal fluctuations in the availability of nitrate (Fig. 3, Supporting Information Fig. S5), ammonium, and phosphate in the soil (Supporting Information Fig. S6, S7). Specifically, the highest concentrations of nitrate and ammonium were observed during spring (March – May) (Supporting Information Table S11). Furthermore, we identified a robust interaction between soil moisture and season with respect to soil nitrate and ammonium levels (Fig 3, Supporting Information Fig. S8, Table S11). Seasonal effects on soil phosphate were not evident, but we did observe a significant positive of soil moisture on phosphate availability (Supporting Information Fig. S8, Table S11, Mean slope = 0.15, 95% CI [0.09, 0.22]). Notably, when assessing these variations along the gradient of tree species richness and among different mycorrhizal types, we found substantial effects only in the case of nitrate (Supporting Information Table S11). Nitrate availability in soil decreased with increasing tree species richness (Fig. 3, Mean slope = −0.09, 95% CI [-0.15, −0.03]). Regarding mycorrhizal types, nitrate availability displayed no consistent pattern, but we did observe interactions between mycorrhizal type and seasonal effects (Fig. 3, Supporting Information Table S11). In stands with EM trees, soil nitrate availability peaked in early spring, surpassing the nitrate levels in AM tree communities, and declined notably during the summer during summer. In AM tree stands, the availability of nitrate reached its peak in May and stayed relatively stable throughout summer. The temporal pattern of further environmental variables can be found in Supporting Information Fig. S9, S10; not included in main analyses because of missing main effects.

**Fig. 3.**
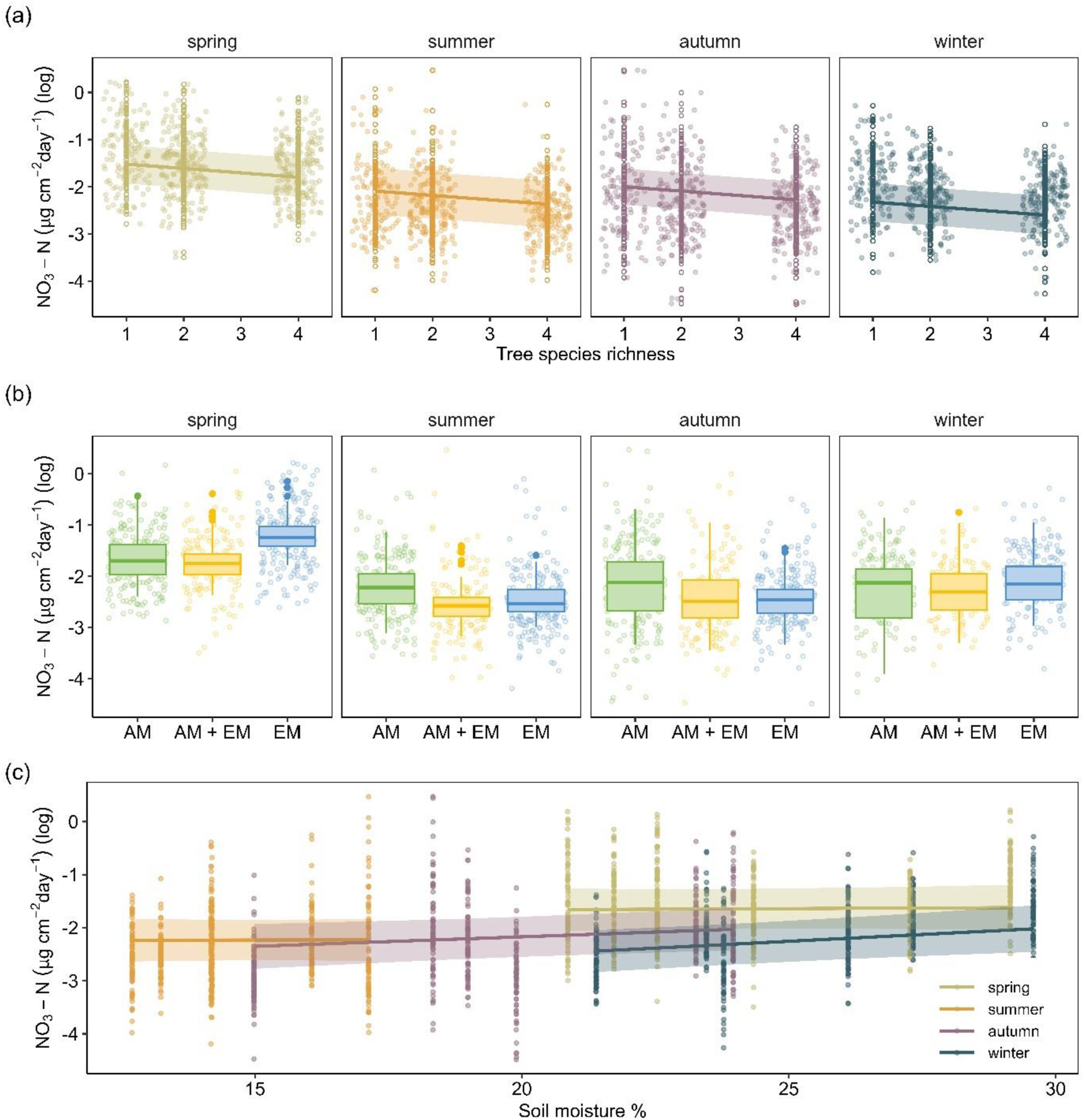
Changes in seasonal availability of nitrate (NO_3_–N) in soil as affected by (a) Tree species richness (one, two, four; Sr), (b) Mycorrhizal type (AM, EM, AM+EM; Myc), and (c) Soil moisture (%). The data points depict the raw observations, regression lines and the shaded bands in panels (a) and (c), and boxplots in panel (b) denote 95% credible intervals, which were derived from predictions generated by a random-walk time series model employing Bayesian approach (see Methods).

### Biodiversity effects

Overall, we found tree species richness to significantly increase net biodiversity effects (sum of complementarity effects and selection effects) for foliar C (*p* = 0.001) and P (*p* <.001) pools, and to marginally increase net biodiversity effects for N pools (*p* = 0.085) (Table 2, Fig. 4; (Supporting Information Table S14). Values were significantly different from zero for all elemental pools in four-species communities (AM, EM, and AM+EM), except for N pools in AM tree stands (Supporting Information Table S12). In addition, tree species richness significantly increased selection effects for all elemental pools (C: *p* <.001, N: *p* = 0.013, P: *p* <.001) and complementarity effects for C pools (*p* = 0.041) and P pools (*p* = 0.007). For all elemental pools in EM tree communities, and N pools in AM tree communities, the positive net biodiversity effects were mostly driven due to strong selection effects, being larger than or similar to the complementarity effects. This is also apparent when comparing selection effects of communities with tree stands of mixed mycorrhizal type (AM+EM) with EM tree strands, where selection effects were higher in the latter for all elemental pools. Contrary, in stands with mixed mycorrhizal types (AM+EM) complementarity effects significantly contributed to overall elemental pools (Supporting Information Table S13). These effects were most pronounced for two-species communities, while in four-species communities complementarity effects and selection effects did not differ strongly from each other. In general, we found no interaction effects (Sr x Myc) for any of the elemental pools. Only for C pools, selection effects had the tendency to be larger in mycorrhizal mixtures with the highest species richness.

**Fig. 4.**
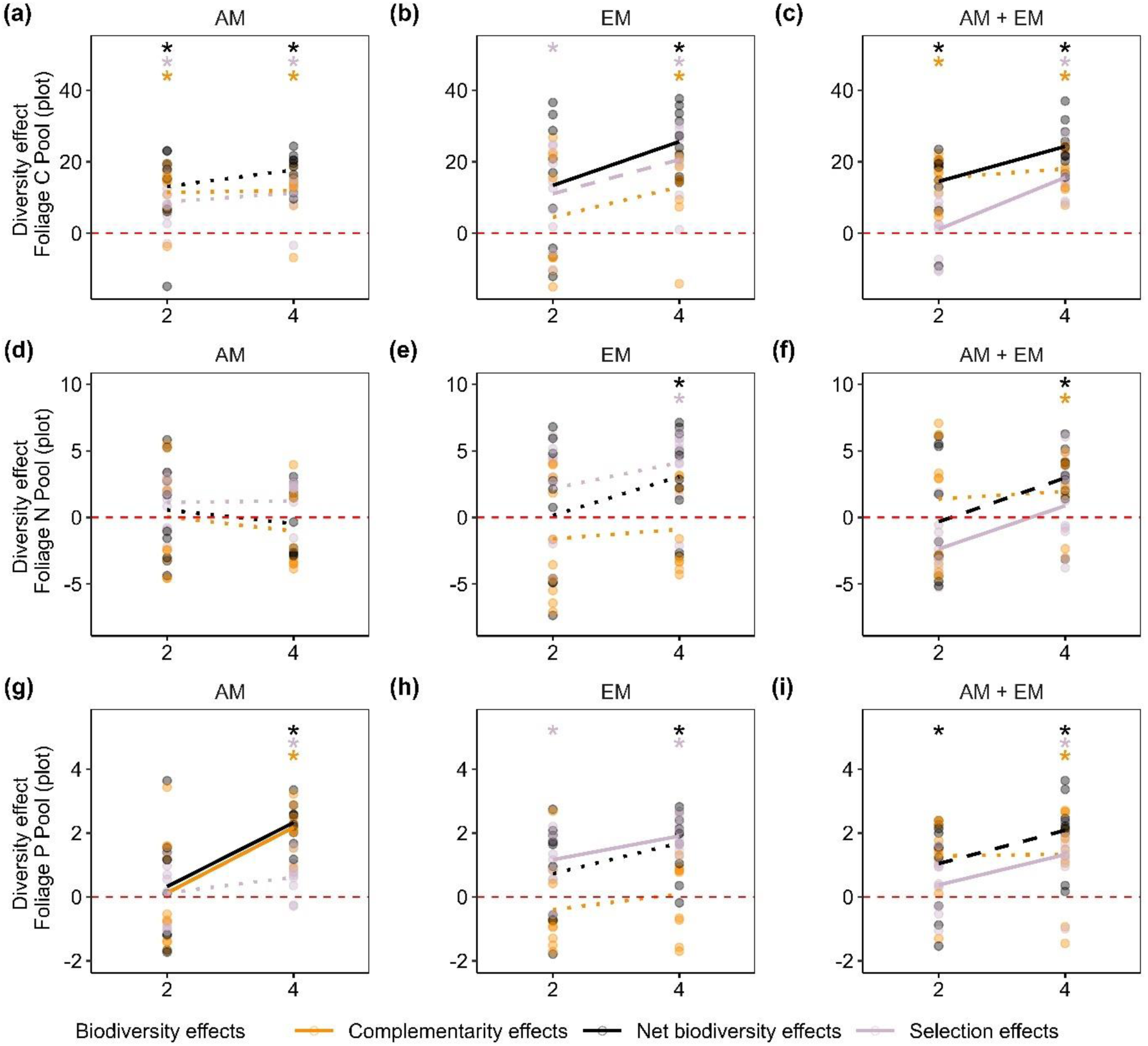
Net biodiversity effects, complementarity effects, and selection effects for (a) carbon (C) pool, (b) nitrogen (N) pool, and (c) phosphorus (P) pool as affected by Tree species richness (2 or 4 tree species) and Mycorrhizal type (AM, EM, AM+EM). Asterisks indicate whether the biodiversity effects were significantly greater than zero (indicating overperformance of two- and four-species tree communities compared to monocultures; detailed results can be found in Supporting Information Table S12). Species specific elemental pools were derived from total pools using species-specific tree volume (see Methods section). Regression lines are based on simple linear regression models (Supporting Information Table S14), whereas solid lines indicate significant relationships between two- and four-species communities (*p* < 0.05), dashed lines marginally significant relationships (0.1 > *p* > 0.05), and dotted lines nonsignificant relationships (*p* > 0.1).

**Table 2.**
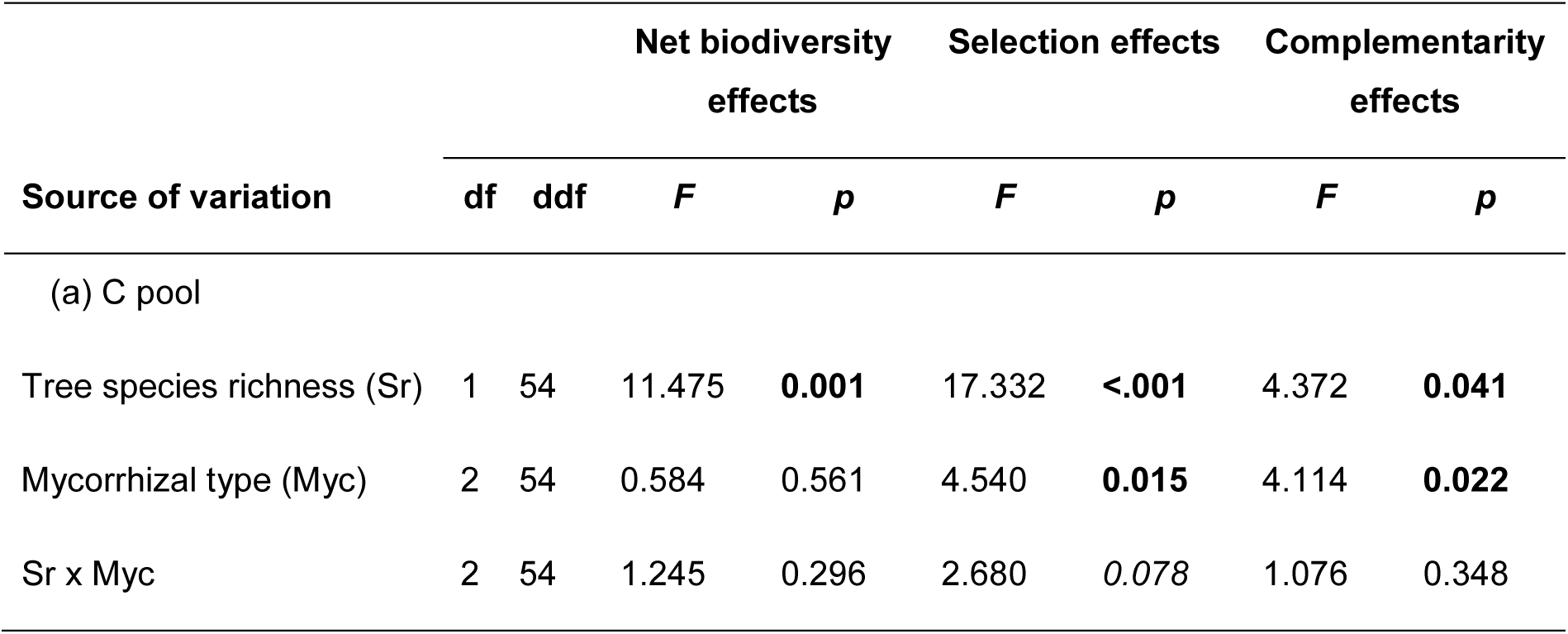
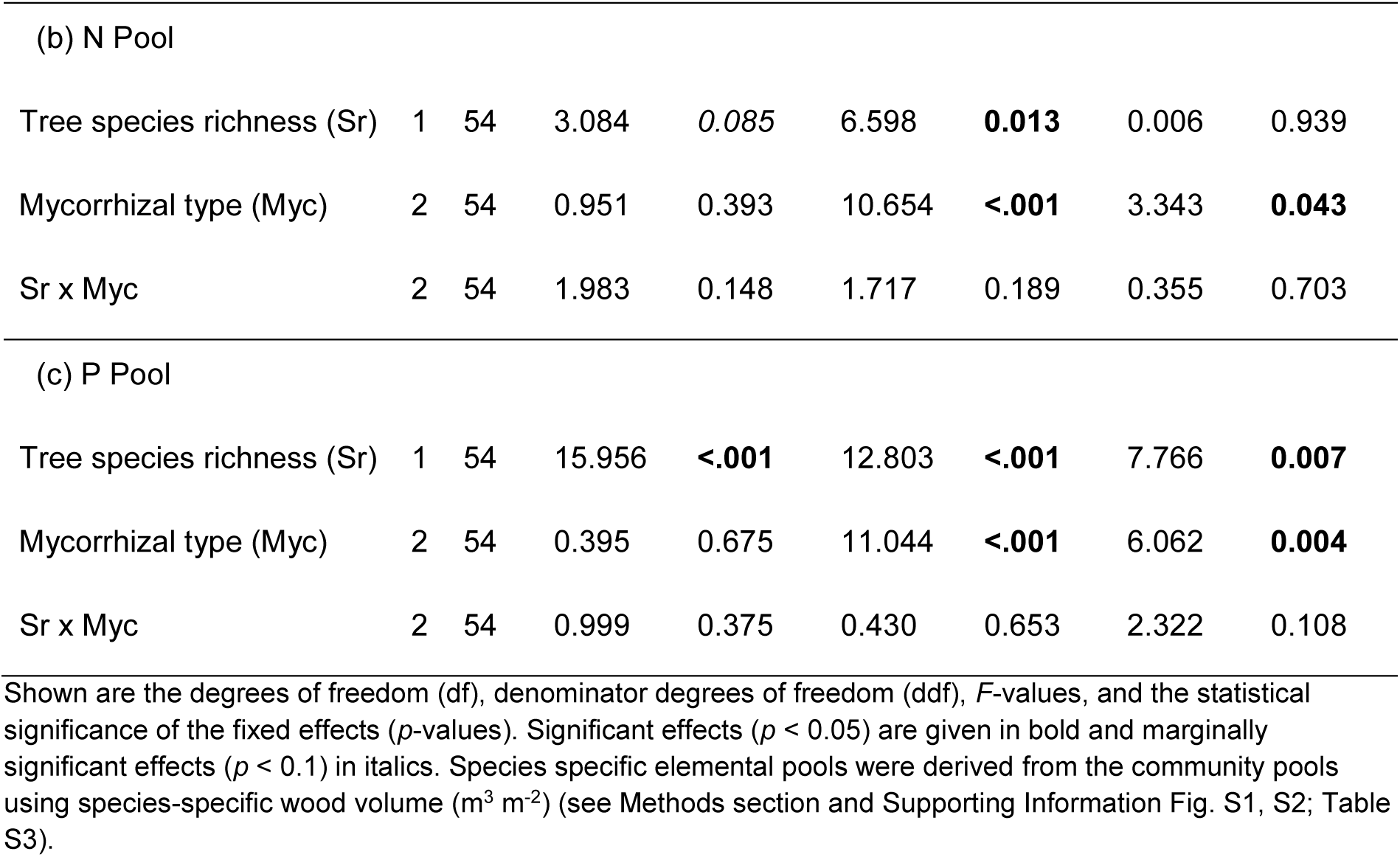
Summary of mixed-effects model analyses testing the effects of tree species richness (two, four; Sr), mycorrhizal type (AM, EM, AM+EM; Myc), and their interactions on net biodiversity effects, selection effects, and complementarity effects based on tree foliage (a) carbon (C) pool, (b) nitrogen (N) pool, and (c) phosphorus (P) pool (n=60).

## Discussion

### Elemental contents and pools are affected by tree species richness and mycorrhizal type

We found significant effects of tree diversity on foliage nutrient contents and pools, whereby the direction of effects differed. While tree diversity had a significantly negative effect on N contents in foliage, P contents and also pools increased with tree species richness. The formation of N pools, however, tended to be increased by tree species richness, contrasting the negative effect on N contents, which may be linked to higher biomass production of more diverse communities as has been shown before both in the MyDiv experiment (Dietrich *et al*., 2022; Ray *et al*., 2023) and in other tree diversity experiments (e.g. Huang *et al*., 2018) and observations (Liang *et al*., 2016; Duffy *et al*., 2017). The higher productivity in more species-rich stands is also supported by the significantly enhanced C pools within these communities. This implies that overall enhanced N uptake at high tree diversity might be masked by an even greater increase in biomass production. Similar patterns were observed in a grassland study where N and P decreased with plant diversity, while C contents remained constant (Guiz *et al*., 2018). Therefore, our first hypothesis (i.e. increased aboveground elemental contents and pools in tree foliage with higher tree diversity) was partly confirmed.

Further, all elemental contents and pools were significantly or at least marginally affected by the mycorrhizal type. Thus, we found greater N contents and pools in EM tree stands compared to AM tree stands, while P contents and pools were significantly enhanced in the latter. The widely reported main roles of mycorrhizal types, being EM fungi providing mainly N to plants and AM fungi supplying plants mainly with P (Smith & Read, 2008; Phillips *et al*., 2013), could therefore also be confirmed for mycorrhizal associations with trees in a young forest plantation (H1). However, in tree stands with both mycorrhizal strategies, we found only additive effects but not an expected elemental pool size that outperforms communities with one mycorrhizal type alone. This is consistent with findings of Dietrich *et al*. (2022) who could not observe any significant overyielding effects of tree productivity in communities of both mycorrhizal types. This is in contrast to our first hypothesis that optimized resource acquisition in mixed stands (Ferlian *et al*., 2018; Eisenhauer *et al*., 2022) will lead to larger elemental pools in foliage.

While aboveground foliage elemental pools and total contents showed significant effects of tree species richness, belowground elemental pools and contents of soil nutrients and microbes were not significantly affected. Only plant-available nutrients showed a clear effect of tree species richness and mycorrhizal type. These limited effects in soils may result from the young age of the tree stands in the MyDiv experiment (seven years since establishment in 2015 until the time point of sampling for this study), given that plant diversity effects on ecosystem functioning have been shown to increase over time in grassland and forest biodiversity experiments (Guerrero-Ramírez *et al*., 2017). Changes in biogeochemical properties may need a longer period of time to establish (Oelmann *et al*., 2011; Lange *et al*., 2023). Further, land-use legacies from former agriculture might still persist and thus it is likely that the trajectory of the microbial soil community development is not be affected by tree species richness in such a short time (Fichtner *et al*., 2014). Missing effects of tree species richness or mycorrhizal type on soil pH and bulk density (Supporting Information Table S4) underpin this assumption. Furthermore, the plastic tarp, initially installed to prevent the growth of weeds that introduce other mycorrhizal types, may have inhibited some processes in nutrient cycling, such as accumulation of organic matter through litter fall and its direct interaction with soil organisms (Berg & McClaugherty, 2014). For example, plants usually modulate their belowground microbiome partly through the quality and composition of litter (Pollierer *et al*., 2007; Prada-Salcedo *et al*., 2022). Therefore, we expect that species richness effects would be more pronounced in case of direct litter input. However, nutrient cycling can be affected by further processes, such as leaching of water-soluble compounds from litter, which plays a significant role in nutrient return to soil (Chapin *et al*., 2012) or the dynamics of root exudates, which were found to be influenced by diversity in tree experiments (Weinhold *et al*., 2022). These processes were not hindered by the tarp. Consequently, we do not anticipate a strong influence of the tarp on our results. In contrast to bulk soil, N and P contents of soil microbial biomass increased with tree species richness, according to our first hypothesis. This observation may indicate that rhizodeposits play a critical role in tree effects on soil food webs (e.g. Pollierer *et al*., 2007) as well as plant diversity effects on soil communities and functions (Lange *et al*., 2015; Eisenhauer *et al*., 2017). While tree diversity had an effect on microbial biomass, we could not find any effects of mycorrhizal type. This is in contrast to further tree diversity studies which found effects of tree diversity or mycorrhizal associations on soil microbial community composition (Singavarapu *et al*., 2021; Ferlian *et al*., 2021).

### Effects of mycorrhizal type on nitrate content are strongly dependent on season and soil moisture

Along with the information on C, N, and P in bulk soil and soil microbial biomass, we assessed plant-available nutrients in soil to underpin soil-related analyses of resource-use complementarity. Tree species richness showed consistent negative effects on soil nutrient availability, while mycorrhizal type showed differing effects on contents of plant-available nutrients in different seasons. Consistent with our second hypothesis, we found that nitrate content decreased significantly with tree species richness. This effect was likely caused by stronger exploitation of soil nitrate due to more complementary resource-use strategies in more diverse tree stands (Ferlian *et al*., 2018; Barry *et al*., 2019).

Effects of mycorrhizal type on nitrate content were strongly dependent on season, showing significantly higher levels of nitrate in EM communities in spring compared to summer and autumn, while nitrate contents in AM communities did not change significantly during these seasons. Soil phosphate and ammonium contents showed pronounced temporal dynamics, but were less affected by the experimental treatments. In general, contents of ammonium and phosphate in soils were low, due to fast immobilization by plants and microorganisms and strong adsorption to soil- and organic colloids (Schachtman *et al*., 1998; Varma *et al*., 2017). These processes may potentially smooth the effects caused by species richness and mycorrhizal types. Contrary, nitrate, as a negatively charged ion, is not easily absorbed by clay particles in soil, and therefore is very mobile and prone to runoff and leaching (Riley *et al*., 2001).

The seasonality of nutrient availability observed is closely linked to the activity of soil organisms and of seasonal patterns of rainfall which is also subject to strong temporal fluctuations (Bardgett & van der Putten, 2014; Baldrian, 2017). Thereby, the seasonally dependent photosynthetic activity of plants and the allocation of photosynthates to the soil play major roles in resource provisioning to soil organisms (Baldrian, 2017). Weather conditions such as temperature, precipitation, and resulting soil moisture (Bardgett & van der Putten, 2014) are important predictors of the activity of soil biodiversity (Bonato Asato *et al*., 2023). In particular, soil water availability has strong effects on recovery and recurrence of soil microbial communities (Placella & Firestone, 2013). Fast responses were observed for nitrifying bacteria (*Nitrobacter* spp., *Nitrospina* spp., and *Nitrospira*) after rewetting of dry soil, followed by significant nitrogen mineralization (Placella & Firestone, 2013). This underlines our findings of high nitrate availability after summer droughts. In addition, studies on litter decay of AM and EM tree communities showed the influence of seasonally variable environmental factors (precipitation and temperature) on mycorrhizal activity (Keller & Phillips, 2018). The reported responses of decay of EM tree litter with higher mean annual precipitation were significantly stronger compared to AM tree litter and also slightly stronger with increases in mean annual temperature (Keller & Phillips, 2018). These findings by Keller & Phillips (2018) may suggest for our observations a higher sensitivity of EM trees towards higher temperatures and moisture and thus, stronger nutrient release from elevated decay of litter.

### Net effects in tree stands of mixed mycorrhizal strategy are driven by complementarity

We found positive net biodiversity effects for C and P pools in foliage. For all elemental pools, net biodiversity effects, selection effects, and complementarity effects increased with tree species richness, except for N pools in AM tree stands. This shows that both, complementarity effects and selection effects, contributed to overall biodiversity effects on nutrient accumulation in foliage in young tree stands. Further, for all elemental pools in four-species communities (AM, EM, AM+EM), except N pools in AM tree stands, net biodiversity effects were significantly different from zero, which indicates overperformance of species-rich communities compared to monocultures (Loreau & Hector, 2001). Similar effects were observed before for other ecosystem functions, such as biomass production in several forest studies (Huang *et al*., 2018; Dietrich *et al*., 2022). For the formation of elemental pools of EM tree stands as well as of N pools of AM tree stands, net biodiversity effects were mostly driven by selection effects, whereas complementarity effects were often lower or even negative. This suggests that the observed larger elemental pools in species-rich tree stands are rather the result of the performance of single dominant tree species (Scherer-Lorenzen *et al*., 2005; Morin *et al*., 2011; Dietrich *et al*., 2022). The observation of high selection effects in EM tree stands, may be attributed to the pronounced nutrient uptake of *Betula pendula* Roth, a fast-growing pioneer tree species (Stark *et al*., 2015), thus contributing a large part to the tree community nutrient pools (Supporting Information Fig. S3). This observation was mainly made for the formation of C, N, and P pools in tree foliage of EM communities with two and four tree species. However, for the generation of foliage P pools in AM four-species communities, complementarity effects were of greater importance. This indicates that the underlying complementarity among trees in the use of P increases P contents and P pools in foliage within higher-diversity communities.

We found that net biodiversity effects tended to be more strongly driven by complementarity effects in mixed tree stands featuring both mycorrhizal types. Therefore, our findings suggest that mixed mycorrhizal strategies may enhance resource partitioning among associated plant hosts and alter aboveground nutrient dynamics in tree foliage (Cheng *et al*., 2016; Ferlian *et al*., 2018; Eisenhauer *et al*., 2022). Although not significant, this was more pronounced for two-species stands compared to four-species stands This result is in line with a recent meta-analysis (Luo *et al*., 2023), which observed highest productivity in stands composed of different mycorrhizal types with equal proportions. Further, that fact that more tree species employ a greater number of resource niches due to increased functional diversity may lead to redundancy in the different roles of mycorrhizal strategies (Luo *et al*., 2023). The third hypothesis (positive effects of high tree species richness and mixed mycorrhizal types on foliage elemental pools can be explained by higher complementarity between tree species and mycorrhizal types) cannot be confirmed.

This study cannot fully explain underlying patterns of positive biodiversity effects, because a direct link between the aboveground and belowground elemental pools was not detected. Elemental assessment of foliage in summer was followed by analyses of soil elements in autumn, which may indicate some temporal mismatch between our measurements. The study of litter chemical composition and quality as well as the derivation of nutrient resorption processes could provide additional insights. Litterfall plays a very important role in nutrient cycling. This will allow to close the gap between the observed patterns in nutrient pools above- and belowground. Further, the analysis of wood and root elemental pools may provide a more comprehensive picture of the above- and belowground dynamics. Our study lacks direct measurements of tree foliage biomass or leaf area index. However, we think that quantification of the SSCI is a very precise proxy to account for these missing measurements. Although we made an effort to include phylogenetic diversity in the study taxa, it is worth noting that four of the five EM tree species belong to a single order, Fagales. Some of our results may therefore reflect the response of a particular lineage (see Koele *et al*., 2012). Clearly, additional research involving a larger group of phylogenetic lineages is warranted.

## Conclusion

We show that tree diversity increases foliage C and P pools, and increases soil nitrate uptake. Both mycorrhizal types thereby contribute differently to aboveground elemental pools, with EM trees forming larger N pools and AM trees forming larger P pools. Elemental pool sizes in tree stands with both mycorrhizal strategies, however, did not exceed those of tree stands with EM or AM alone. However, the analysis of biodiversity effects indicates that resource-use complementarity affects the resource uptake and aboveground nutrient storage in foliage more strongly in tree communities with both mycorrhizal types. Our findings thus emphasize the importance of using forest species with diverse mycorrhizal strategies during restoration for achieving a more complete use of available resources and thus to deliver more multifunctional forests. Given the prevalence of impoverished forests due to anthropogenic impact, and the need to implement effective restoration programs, our results are of broad significance.

## Acknowledgements

We thank Julius Quosh for collecting the dendrometric data and Romy Zeiss, Alla Kavtea, and Tom Künne for collecting the time series data on plant available nutrients (IEMs). Further we thank Victoria Ochoa for the chemical analysis of IEMs and Ines Hilke for the measurements of C, N, and P in a the MPI-BGC Jena. We acknowledge funding by the Deutsche Forschungsgemeinschaft (German Centre for Integrative Biodiversity Research, FZT118; and Gottfried Wilhelm Leibniz Prize, Ei 862/29-1). FTM acknowledges funding from the European Research Council (ERC Grant agreement 647038 [BIODESERT]) and Generalitat Valenciana (CIDEGENT/2018/041).

## Competing Interests

The authors declare no competing interests.

## Author contributions

N.E. and O.F. designed and established the experiment.

N.E. acquired the funds for this project.

E.Bö., J.L., F.T.M, E.B. and T.R. acquired the data.

E.Bö. analyzed the data and created the figures and wrote the manuscript.

Y.H. analyzed the time series data on plant available nutrients.

All authors significantly revised the manuscript and approved it for submission.

## Data availability

The data that support the findings of this study are deposited in MyDiv database (https://mydivdata.idiv.de/) and will be published after acceptance of the manuscript. Access codes will then be made available.

## Supporting Information

Additional Supporting Information may be found online in the Supporting Information section at the end of the article.

**Fig. S1** Wood volume for tree communities of AM, EM, AM+EM trees with one, two, four tree species.

**Fig. S2** Species specific wood volume for tree communities of AM, EM, AM+EM trees with one, two, four tree species.

**Fig. S3** Species specific foliage elemental pools for tree communities of AM, EM, AM+EM trees with one, two, four tree species.

**Fig. S4** Soil and soil microbial elemental pools for tree communities of AM, EM, AM+EM trees with one, two, four tree species.

**Fig. S5** Temporal variability of nitrate availability in soil over the period of two years.

**Fig. S6** Temporal variability of ammonium availability in soil over the period of two years.

**Fig. S7** Temporal variability of phosphate availability in soil over the period of two years.

**Fig. S8** Seasonal availability of ammonium and phosphate as affected by soil moisture.

**Fig. S9** Temporal variation of soil temperature over the period of two years.

**Fig. S10** Temporal variation of air temperature and air humidity over the period of two years.

**Table S1** Plot specific information of MyDiv experiment.

**Table S2** Summary Tukey HSD analysis of wood volume among Mycorrhizal types.

**Table S3** Summary mixed-effects model analyses testing the effects of Tree species richness and Mycorrhizal type on wood volume.

**Table S4** Summary mixed-effects model analyses testing the effects of Tree species richness and Mycorrhizal type on soil pH and soil bulk density.

**Table S5** Summary mixed-effects model analyses testing the effects of Tree species richness, Mycorrhizal type, and their interaction of elemental contents in foliage, soil, and soil microbial biomass.

**Table S6** Summary mixed-effects model analyses testing the effects of Tree species richness, Mycorrhizal type, and their interaction of elemental pools in foliage, soil, and soil microbial biomass.

**Table S7** Summary Tukey HSD analysis of elemental contents in foliage, soil, and soil microbial biomass among Mycorrhizal types.

**Table S8** Summary Tukey HSD analysis of elemental pools in foliage, soil, and soil microbial biomass among Mycorrhizal types.

**Table S9** Summary simple linear regression analyses of elemental contents in foliage, soil, and soil microbial biomass.

**Table S10** Summary simple linear regression analyses of elemental pools in foliage, soil, and soil microbial biomass.

**Table S11** Summary of RW1 model using Bayesian statistics testing the effects of Tree species richness, Mycorrhizal type, Season, and Soil moisture nitrate ammonium phosphate availability.

**Table S12** Summary one-tailed t-test to test whether biodiversity effects are significantly different from zero.

**Table S13** Summary Tukey HSD analysis testing the difference in biodiversity effects (net biodiversity effects, selection effects, complementarity effects) among Mycorrhizal types.

**Table S14** Summary of simple linear regression analyses of biodiversity effects of mycorrhizal tree communities.

**Table S15** Percentage changes of elemental contents in foliage, soil, and soil microbial biomass between levels of Tree species richness.

**Table S16** Percentage changes of elemental pools in foliage, soil, and soil microbial biomass between levels of Tree species richness.

**Table S17** Percentage changes of elemental contents in foliage, soil, and soil microbial biomass between levels of Mycorrhizal type.

**Table S18** Percentage changes of elemental pools in foliage, soil, and soil microbial biomass between levels of Mycorrhizal type.

## Notes

### Competing Interest Statement

The authors have declared no competing interest.

