## Supporting Information for "Mycorrhizal fungi modulate tree diversity effects on nutrient dynamics"

New Phytologist

**Article acceptance date**: [...]

**The following Supporting Information is available for this article**:

**Fig. S1** Wood volume for tree communities of AM, EM, AM+EM trees with one, two, four tree species.

**Fig. S2** Species specific wood volume for tree communities of AM, EM, AM+EM trees with one, two, four tree species.

**Fig. S3** Species specific foliage elemental pools for tree communities of AM, EM, AM+EM trees with one, two, four tree species.

**Fig. S4** Soil and soil microbial elemental pools for tree communities of AM, EM, AM+EM trees with one, two, four tree species.

**Fig. S5** Temporal variability of nitrate availability in soil over the period of two years.

**Fig. S6** Temporal variability of ammonium availability in soil over the period of two years.

**Fig. S7** Temporal variability of phosphate availability in soil over the period of two years.

**Fig. S8** Seasonal availability of ammonium and phosphate as affected by soil moisture.

**Fig. S9** Temporal variation of soil temperature over the period of two years.

**Fig. S10** Temporal variation of air temperature and air humidity over the period of two years.

**Table S1** Plot specific information of MyDiv experiment.

**Table S2** Summary Tukey HSD analysis of wood volume among Mycorrhizal types.

**Table S3** Summary mixed-effects model analyses testing the effects of Tree species richness and Mycorrhizal type on wood volume.

**Table S4** Summary mixed-effects model analyses testing the effects of Tree species richness and Mycorrhizal type on soil pH and soil bulk density.

**Table S5** Summary mixed-effects model analyses testing the effects of Tree species richness, Mycorrhizal type, and their interaction of elemental contents in foliage, soil, and soil microbial biomass.

**Table S6** Summary mixed-effects model analyses testing the effects of Tree species richness, Mycorrhizal type, and their interaction of elemental pools in foliage, soil, and soil microbial biomass.

**Table S7** Summary Tukey HSD analysis of elemental contents in foliage, soil, and soil microbial biomass among Mycorrhizal types.

**Table S8** Summary Tukey HSD analysis of elemental pools in foliage, soil, and soil microbial biomass among Mycorrhizal types.

**Table S9** Summary simple linear regression analyses of elemental contents in foliage, soil, and soil microbial biomass.

**Table S10** Summary simple linear regression analyses of elemental pools in foliage, soil, and soil microbial biomass.

**Table S11** Summary of RW1 model using Bayesian statistics testing the effects of Tree species richness, Mycorrhizal type, Season, and Soil moisture nitrate ammonium phosphate availability.

**Table S12** Summary one-tailed t-test to test whether biodiversity effects are significantly different from zero.

**Table S13** Summary Tukey HSD analysis testing the difference in biodiversity effects (net biodiversity effects, selection effects, complementarity effects) among Mycorrhizal types.

**Table S14** Summary of simple linear regression analyses of biodiversity effects of mycorrhizal tree communities.

**Table S15** Percentage changes of elemental contents in foliage, soil, and soil microbial biomass between levels of Tree species richness.

**Table S16** Percentage changes of elemental pools in foliage, soil, and soil microbial biomass between levels of Tree species richness.

**Table S17** Percentage changes of elemental contents in foliage, soil, and soil microbial biomass between levels of Mycorrhizal type.

**Table S18** Percentage changes of elemental pools in foliage, soil, and soil microbial biomass between levels of Mycorrhizal type.

Fig. S1 Wood volume of trees ($m^3$ $m^{-2}$) as a function of Tree species richness (one, two, four; Sr) for communities containing arbuscular mycorrhizal tree species (AM), ectomycorrhizal tree species (EM), or both (AM + EM) tree species (Myc). Each dot represents a tree community, and colors indicate different Mycorrhizal types. Regression lines are based on mixed-effect models (predicted means). Solid lines indicate statistically significant relationships ($p$ < 0.05), dashed lines marginally significant relationships (0.1 > $p$ > 0.05), and dotted lines non-significant relationships ($p$ > 0.1). Statistical significance of main effects is indicated in each panel (n.s., $p$ > 0.05; * $p$ < 0.05; ** $p$ <0.01; *** $p$ <0.001).

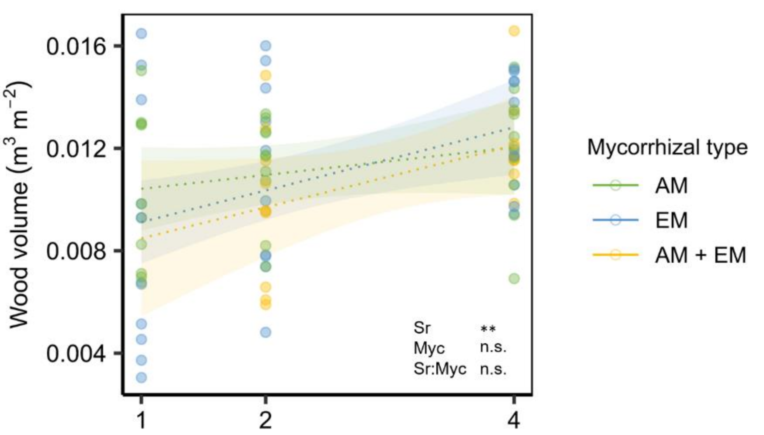


Fig. S2 Wood volume for each tree species, indicated by different colors, according to Mycorrhizal type (AM, EM, AM+EM; Myc) and Tree species richness (one, two, four; Sr). Lower number of individuals in mixtures was corrected by multiplying basal area with species richness.

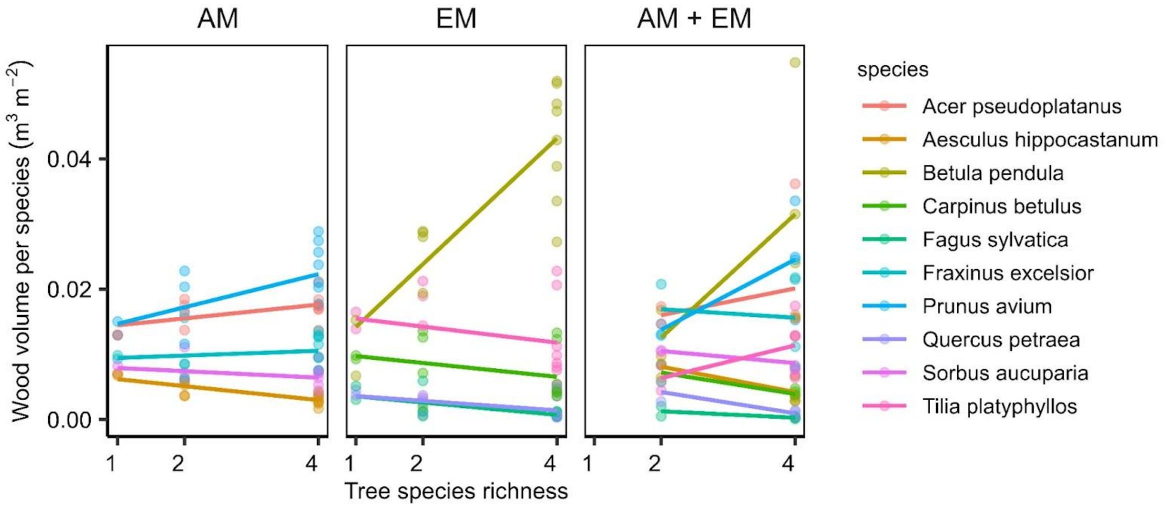

Fig. S3 (a) Carbon (C), (b) nitrogen (N) and (c) phosphorus (P) pools of foliage for each tree species, indicated by different colors, according to Mycorrhizal type (AM, EM, AM+EM; Myc) and Tree species richness (one, two, four; Sr). Lower number of individuals in mixtures was corrected by multiplying basal area with species richness.

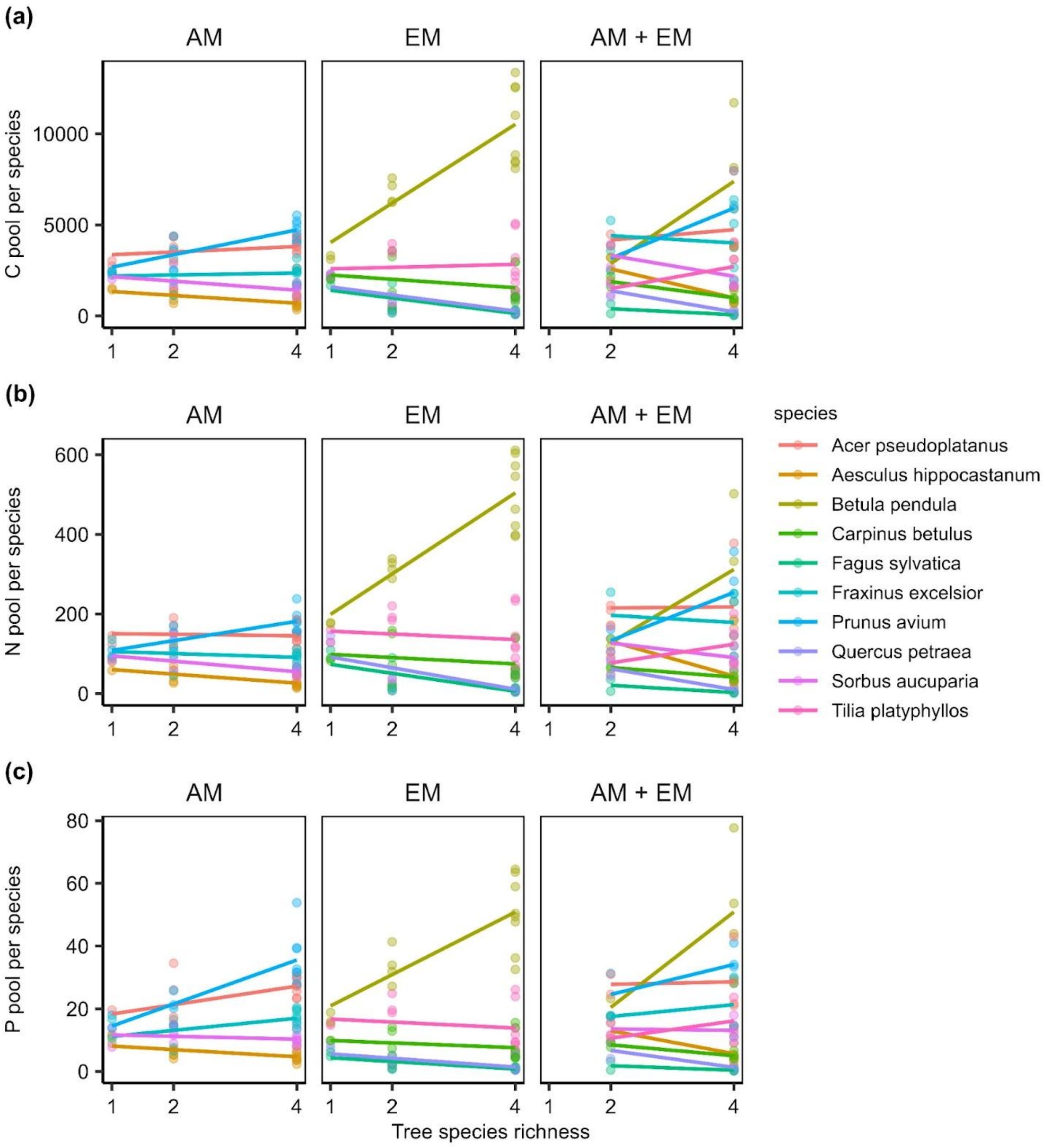

Fig. S4 Carbon (C), nitrogen (N) and phosphorus (P) pools of soil (a-c) and microbial biomass (d-f) as a function of Tree species richness (one, two, four; Sr) for communities containing arbuscular mycorrhizal tree species (AM), ectomycorrhizal tree species (EM), or both (AM + EM) tree species (Myc). Each dot represents a tree community, and colors indicate different Mycorrhizal types. Regression lines are based on mixed-effect models (predicted means). Solid lines indicate statistically significant relationships ($p < 0.05$), dashed lines marginally significant relationships ($0.1 > p > 0.05$), and dotted lines non-significant relationships ($p > 0.1$). No statistically significant main effects found.

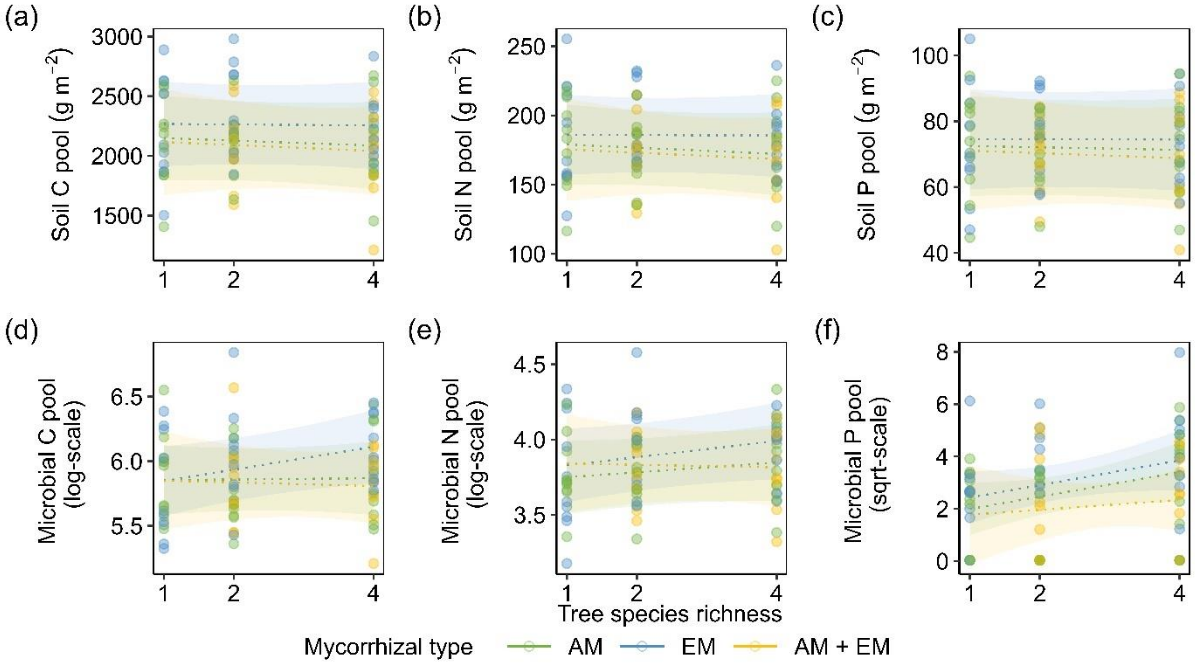

Fig. S5 Availability of nitrate ($NO_3$-N) in soil over the period of two years (April 2019 - March 2021) as affected by (a) Tree species richness (one, two, four; Sr) and (b) Mycorrhizal type (AM, EM, AM+EM).

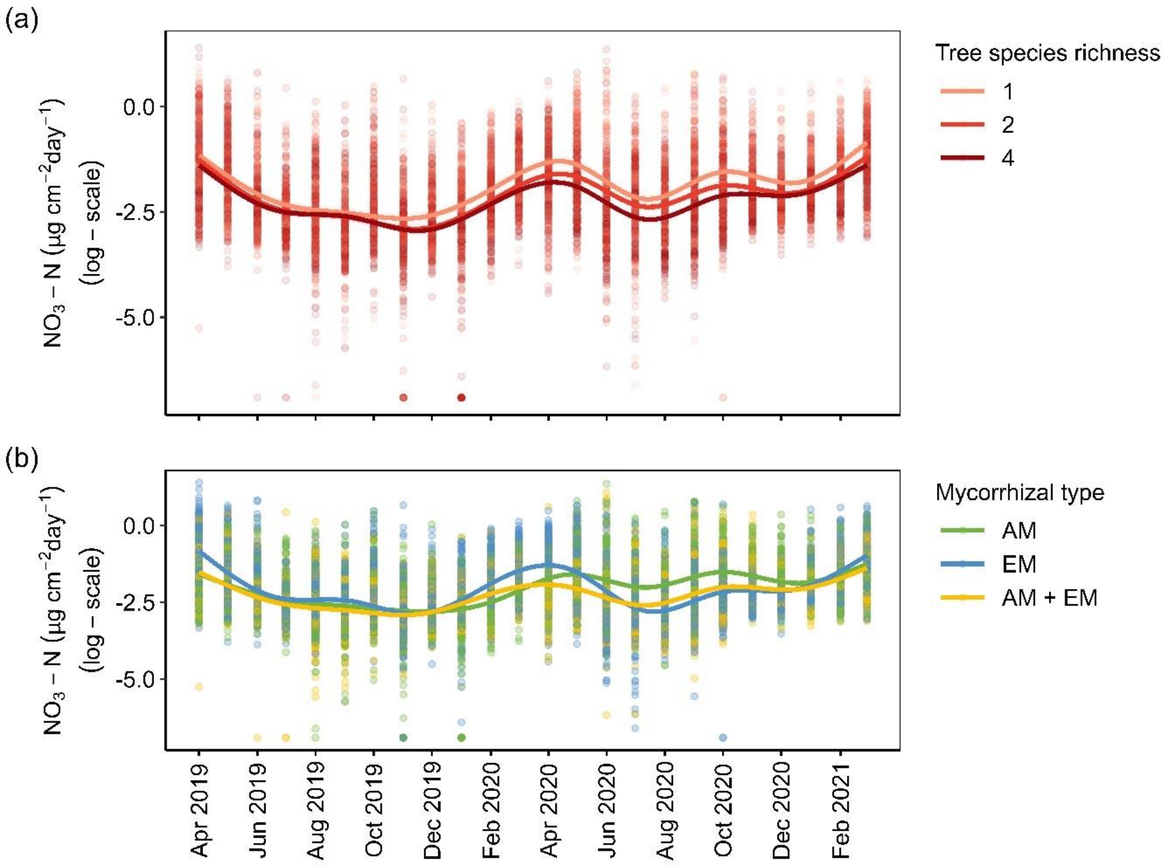

Fig. S6 Availability of ammonium ($NH_4$-N) in soil over the period of two years (April 2019 - March 2021) as affected by (a) Tree species richness (one, two, four; Sr) and (b) Mycorrhizal type (AM, EM, AM+EM). No statistically significant effects of both treatments or their interaction on ammonium found.

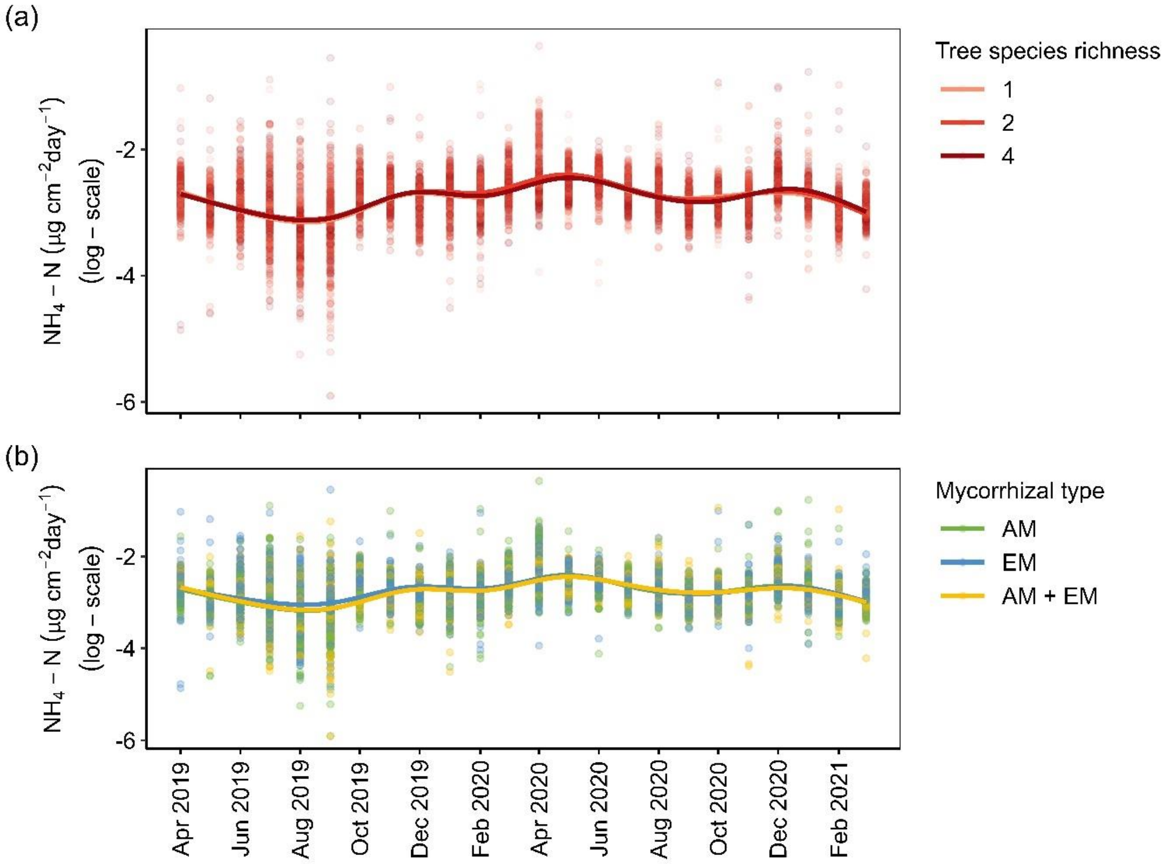

Fig. S7 Availability of phosphate ($PO_4$-P) in soil over the period of two years (April 2019 - March 2021) as affected by (a) Tree species richness (one, two, four; Sr) and (b) Mycorrhizal type (AM, EM, AM+EM). No statistically significant effects of both treatments or their interaction on phosphate found.

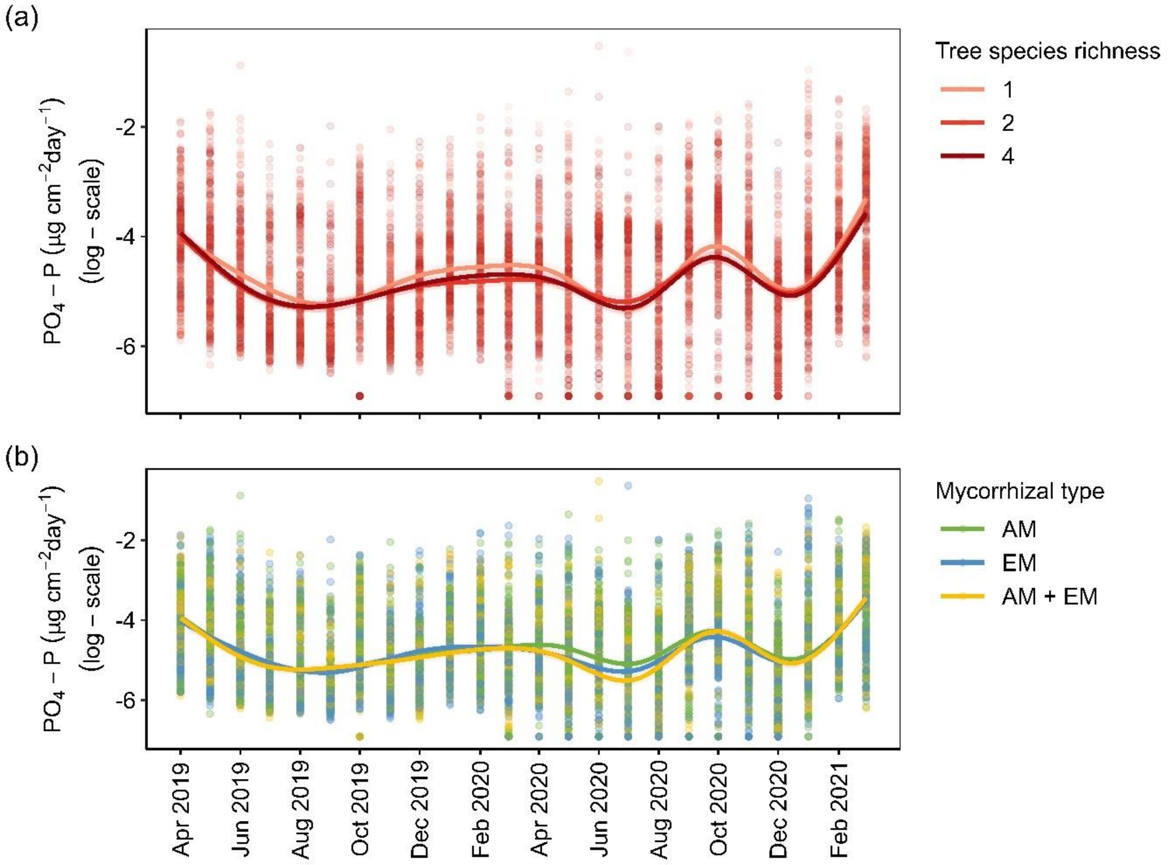

Fig. S8 Seasonal availability of (a) ammonium ($NH_4$-N), and (b) phosphate ($PO_4$-P) as affected by soil moisture (%). Results can be found in Supporting Information Table S….

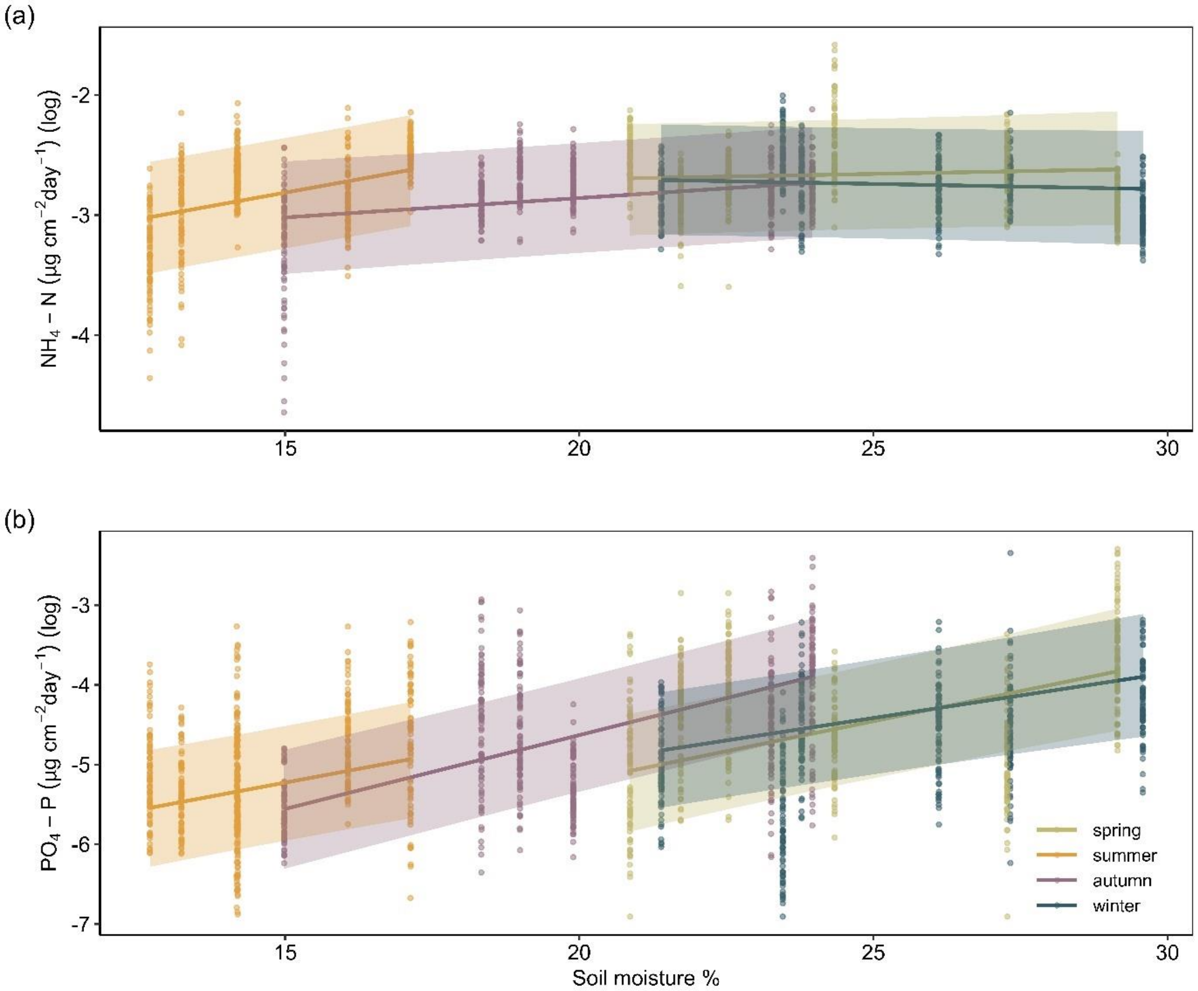


Fig. S9 Temporal variation in soil temperature (10 cm depth) over the period of two years (April 2019 - March 2021). No statistically significant effect of soil temperature on plant available nutrients found.

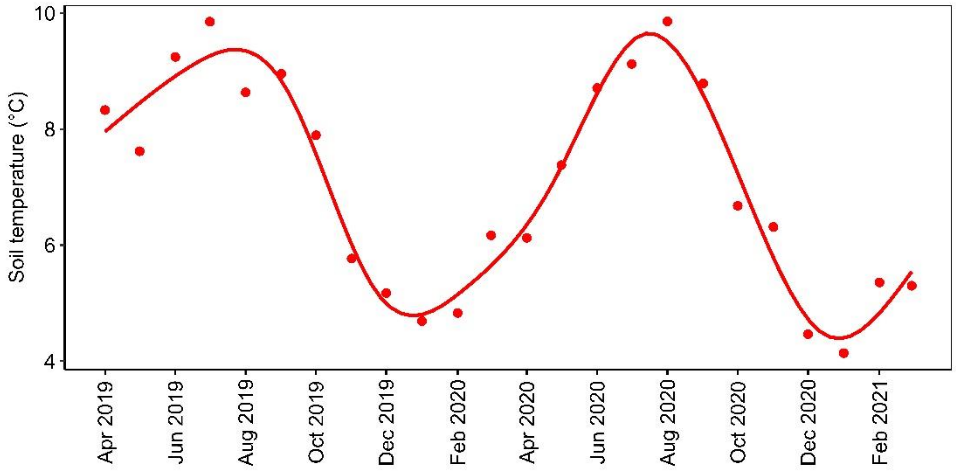

Fig. S10 Temporal variation in air temperature (°C) and air humidity (%) over the period of two years (April 2019 - March 2021). No statistically significant effects of both parameters on plant available nutrients found.

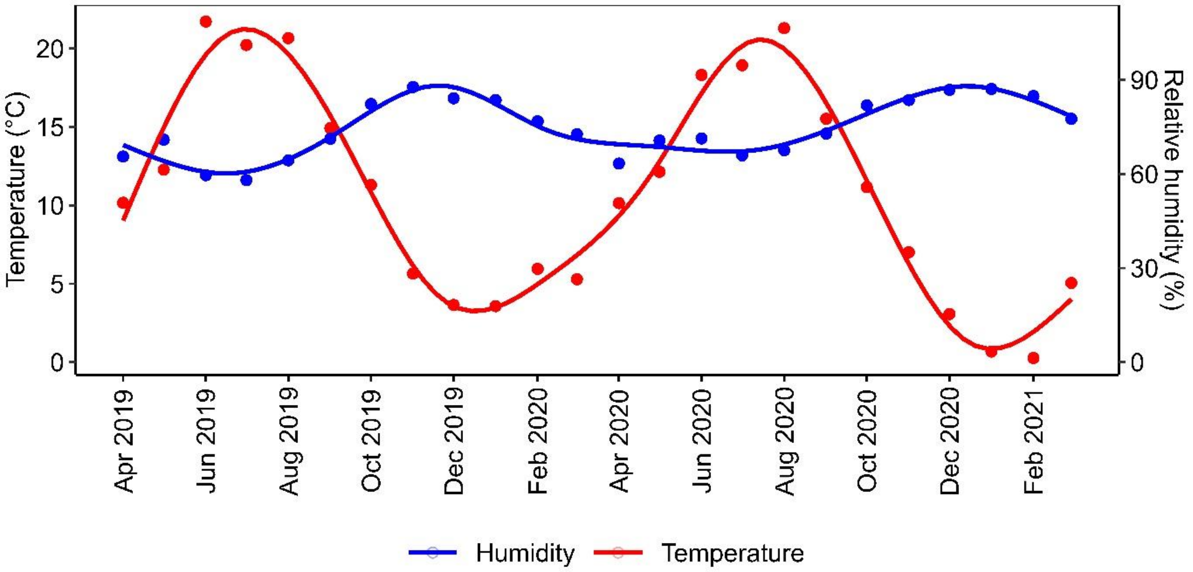

Table S1 MyDiv experimental design with block (one, two), plot ID (1 to 80), and main treatments Tree species richness (one, two, four; Sr), Mycorrhizal type (AM, EM, AM+EM; Myc), and the species composition (Ac - *Acer pseudoplatanus* L., Ae - *Aesculus hippocastanum* L., Fr - *Fraxinus excelsior* L., Pr - *Prunus avium* L., So - *Sorbus aucuparia* L., Be - *Betula pendula* Roth, Ca - *Carpinus betulus* L., Fa - *Fagus sylvatica* L., Qu - *Quercus petraea* (Matt.) Liebl., Ti - *Tilia platyphyllos* Scop.).

| **Block** | **Plot ID** | **Tree species richness** | **Mycorrhizal type** | **Species composition** |
|---|---|---|---|---|
| 1 | 1 | 2 | AM+EM | Pr\|Ti |
| 1 | 2 | 1 | EM | Fa |
| 1 | 3 | 2 | AM | Fr\|So |
| 1 | 4 | 1 | EM | Qu |
| 1 | 5 | 2 | EM | Be\|Qu |
| 1 | 6 | 4 | AM | Ac\|Ae\|Fr\|So |
| 1 | 7 | 4 | EM | Be\|Ca\|Qu\|Ti |
| 1 | 8 | 1 | AM | Fr |
| 1 | 9 | 2 | AM | Ac\|Pr |
| 1 | 10 | 4 | AM | Ae\|Fr\|Pr\|So |
| 1 | 11 | 2 | EM | Qu\|Ti |
| 1 | 12 | 4 | EM | Be\|Ca\|Fa\|Ti |
| 1 | 13 | 2 | EM | Ca\|Fa |
| 1 | 14 | 4 | AM+EM | Pr\|So\|Be\|Qu |
| 1 | 15 | 2 | AM+EM | Fr\|Ca |
| 1 | 16 | 4 | AM+EM | Ac\|Ae\|Be\|Ca |
| 1 | 17 | 1 | AM | So |
| 1 | 18 | 2 | AM+EM | Ac\|Be |
| 1 | 19 | 1 | AM | Ac |
| 1 | 20 | 1 | EM | Ti |
| 1 | 21 | 4 | AM | Ac\|Fr\|Pr\|So |
| 1 | 22 | 1 | EM | Ca |
| 1 | 23 | 2 | AM+EM | Ae\|Fa |
| 1 | 24 | 4 | AM+EM | Fr\|So\|Ca\|Fa |
| 1 | 25 | 2 | AM | Pr\|So |
| 1 | 26 | 4 | AM | Ac\|Ae\|Pr\|So |
| 1 | 27 | 1 | EM | Be |
| 1 | 28 | 4 | AM+EM | Ae\|Fr\|Qu\|Ti |
| 1 | 29 | 2 | EM | Fa\|Ti |
| 1 | 30 | 4 | EM | Be\|Fa\|Qu\|Ti |
| 1 | 31 | 1 | AM | Pr |
| 1 | 32 | 4 | EM | Be\|Ca\|Fa\|Qu |
| 1 | 33 | 1 | AM | Ae |
| 1 | 34 | 4 | EM | Ca\|Fa\|Qu\|Ti |
| 1 | 35 | 2 | AM | Ac\|Ae |
| 1 | 36 | 4 | AM+EM | Ac\|Pr\|Fa\|Ti |
| 1 | 37 | 2 | EM | Be\|Ca |
| 1 | 38 | 2 | AM+EM | So\|Qu |
| 1 | 39 | 2 | AM | Ae\|Fr |
| 1 | 40 | 4 | AM | Ac\|Ae\|Fr\|Pr |
| 2 | 41 | 2 | AM+EM | Pr\|Be |

| | | | | |
|---|---|---|---|---|
| 2 | 42 | 4 | AM+EM | Fr\|Pr\|Fa\|Qu |
| 2 | 43 | 2 | EM | Be\|Ti |
| 2 | 44 | 4 | AM | Ae\|Fr\|Pr\|So |
| 2 | 45 | 4 | EM | Be\|Ca\|Qu\|Ti |
| 2 | 46 | 1 | AM | Ac |
| 2 | 47 | 1 | EM | Qu |
| 2 | 48 | 4 | EM | Be\|Ca\|Fa\|Ti |
| 2 | 49 | 4 | AM+EM | Ae\|So\|Be\|Fa |
| 2 | 50 | 1 | AM | Ae |
| 2 | 51 | 2 | EM | Ca\|Qu |
| 2 | 52 | 2 | AM | Ac\|Fr |
| 2 | 53 | 4 | AM | Ac\|Ae\|Fr\|Pr |
| 2 | 54 | 2 | AM | Ac\|So |
| 2 | 55 | 4 | AM+EM | Ae\|Pr\|Ca\|Ti |
| 2 | 56 | 1 | EM | Fa |
| 2 | 57 | 2 | AM+EM | Ae\|Qu |
| 2 | 58 | 4 | EM | Ca\|Fa\|Qu\|Ti |
| 2 | 59 | 4 | AM | Ac\|Ae\|Fr\|So |
| 2 | 60 | 4 | AM+EM | Ac\|Fr\|Be\|Ti |
| 2 | 61 | 1 | AM | Fr |
| 2 | 62 | 2 | EM | Ca\|Ti |
| 2 | 63 | 2 | EM | Be\|Fa |
| 2 | 64 | 2 | AM | Ae\|Pr |
| 2 | 65 | 2 | AM+EM | So\|Ca |
| 2 | 66 | 4 | AM | Ac\|Fr\|Pr\|So |
| 2 | 67 | 2 | AM+EM | Fr\|Fa |
| 2 | 68 | 1 | AM | So |
| 2 | 69 | 2 | AM | Fr\|Pr |
| 2 | 70 | 1 | EM | Ti |
| 2 | 71 | 4 | EM | Be\|Ca\|Fa\|Qu |
| 2 | 72 | 1 | EM | Ca |
| 2 | 73 | 2 | AM | Ae\|So |
| 2 | 74 | 4 | AM | Ac\|Ae\|Pr\|So |
| 2 | 75 | 4 | AM+EM | Ac\|So\|Ca\|Qu |
| 2 | 76 | 1 | EM | Be |
| 2 | 77 | 1 | AM | Pr |
| 2 | 78 | 2 | AM+EM | Ac\|Ti |
| 2 | 79 | 4 | EM | Be\|Fa\|Qu\|Ti |
| 2 | 80 | 2 | EM | Fa\|Qu |

Table S2 Summary of Tukey HSD analysis testing the difference in wood volume ($m^3$ $m^{-2}$) among Mycorrhizal types (AM, EM, AM+EM, Myc) (n=80).

| | | **Wood volume** | |
|---|---|---|---|
| **Source of variation** | **df** | ***t*** | ***p*** |
| AM vs. EM | 73 | 0.327 | 0.943 |
| AM vs. AM+EM | 73 | 0.980 | 0.592 |
| EM vs. AM+EM | 73 | 0.706 | 0.761 |

Table S3 Summary of mixed-effect model analyses (Type-I Sum of Squares) testing the effects of Tree species richness (one, two, four; Sr), Mycorrhizal type (AM, EM, AM+EM; Myc), and their interactions on wood volume ($m^3$ $m^{-2}$) (n=80).

| | | | **Wood volume** | |
|---|---|---|---|---|
| **Source of variation** | **df** | **ddf** | ***F*** | ***p*** |
| Tree species richness (Sr) | 1 | 73 | 10.174 | **0.002** |
| Mycorrhizal type (Myc) | 2 | 73 | 0.461 | 0.633 |
| Sr x Myc | 2 | 73 | 0.695 | 0.502 |

Shown are the degrees of freedom (df), denominator degrees of freedom (ddf), *F*-values, and the statistical significance of the fixed effects (*p*-values). Significant effects ($p < 0.05$) are given in bold and marginally significant effects ($p < 0.1$) in italics.

Table S4 Summary of mixed-effects model analyses (Type-I Sum of Squares) testing the effects of Tree species richness (one, two, four; Sr), Mycorrhizal type (AM, EM, AM+EM; Myc), and their interactions on soil pH and soil bulk density (n=80).

| | | | **pH** | | **Soil bulk density** | |
|---|---|---|---|---|---|---|
| **Source of variation** | **df** | **ddf** | ***F*** | ***p*** | ***F*** | ***p*** |
| Tree species richness (Sr) | 1 | 73 | 0.052 | 0.821 | 0.983 | 0.325 |
| Mycorrhizal type (Myc) | 2 | 73 | 0.553 | 0.577 | 1.381 | 0.258 |
| Sr x Myc | 2 | 73 | 0.159 | 0.853 | 0.195 | 0.823 |

Shown are the degrees of freedom (df), denominator degrees of freedom (ddf), *F*-values, and the statistical significance of the fixed effects (*p*-values). Significant effects ($p < 0.05$) are given in bold and marginally significant effects ($p < 0.1$) in italics.

Table S5 Summary of mixed-effects model analyses (Type-1 Sum of Squares) testing the effects of Tree species richness (one, two, four; Sr), Mycorrhizal type (AM, EM, AM+EM; Myc), and their interactions on (a) foliage, (b) soil, and (c) microbial biomass carbon (C), nitrogen (N), and phosphorus (P) content (g $kg^{-1}$ for foliage, soil; μg $kg^{-1}$ for microbial biomass) (n=80).

| | | | **C content** | | **N content** | | **P content** | |
|---|---|---|---|---|---|---|---|---|
| **Source of variation** | **df** | **ddf** | ***F*** | ***p*** | ***F*** | ***p*** | ***F*** | ***p*** |
| (a) Foliage | | | | | | | | |
| Tree species richness (Sr) | 1 | 73 | 0.606 | 0.439 | 12.467 | **<.001** | 5.941 | **0.017** |
| Mycorrhizal type (Myc) | 2 | 73 | 27.434 | **<.001** | 27.279 | **<.001** | 7.962 | **0.001** |
| Sr x Myc | 2 | 73 | 0.275 | 0.760 | 0.376 | 0.688 | 1.162 | 0.319 |
| (b) Soil | | | | | | | | |
| Tree species richness (Sr) | 1 | 73 | 0.008 | 0.928 | 0.001 | 0.979 | 0.275 | 0.602 |
| Mycorrhizal type (Myc) | 2 | 73 | 0.852 | 0.431 | 0.523 | 0.595 | 0.311 | 0.734 |
| Sr x Myc | 2 | 73 | 0.114 | 0.892 | 0.082 | 0.922 | 0.327 | 0.723 |
| (c) Microbial biomass | | | | | | | | |
| Species richness (Sr) | 1 | 73 | 2.247 | 0.138 | 4.782 | **0.032** | 6.754 | **0.011** |
| Mycorrhizal type (Myc) | 2 | 73 | 0.838 | 0.437 | 0.637 | 0.531 | 1.849 | 0.165 |
| Sr x Myc | 2 | 73 | 0.961 | 0.387 | 0.387 | 0.681 | 0.555 | 0.577 |

Shown are the degrees of freedom (df), denominator degrees of freedom (ddf), *F*-values, and the statistical significance of the fixed effects (*p*-values). Significant effects ($p < 0.05$) are given in bold and marginally significant effects ($p < 0.1$) in italics.

Table S6 Summary of mixed-effects model analyses (Type-I Sum of Squares) testing the effects of Tree species richness (one, two, four; Sr), Mycorrhizal type (AM, EM, AM+EM; Myc), and their interactions on (a) foliage, (b) soil, and (c) microbial biomass carbon (C), nitrogen (N) and phosphorus (P) pools. Elemental pools in foliage represent the product of the elemental concentration with the Stand Structural Complexity Index (SSCI) (n=80).

| | | | **C Pool** | | **N Pool** | | **P Pool** | |
|---|---|---|---|---|---|---|---|---|
| **Source of variation** | **df** | **ddf** | ***F*** | ***p*** | ***F*** | ***p*** | ***F*** | ***p*** |
| (a) Foliage | | | | | | | | |
| Tree species richness (Sr) | 1 | 73 | 15.265 | **<.001** | 2.111 | 0.151 | 16.746 | **<.001** |
| Mycorrhizal type (Myc) | 2 | 73 | 1.677 | 0.194 | 11.416 | **<.001** | 2.818 | *0.066* |
| Sr x Myc | 2 | 73 | 0.883 | 0.418 | 1.005 | 0.371 | 0.353 | 0.704 |
| (b) Soil | | | | | | | | |
| Tree species richness (Sr) | 1 | 73 | 0.695 | 0.407 | 0.738 | 0.393 | 0.303 | 0.584 |
| Mycorrhizal type (Myc) | 2 | 73 | 2.383 | 0.099 | 1.720 | 0.186 | 0.958 | 0.388 |
| Sr x Myc | 2 | 73 | 0.045 | 0.956 | 0.081 | 0.922 | 0.031 | 0.971 |
| (c) Soil microbial biomass | | | | | | | | |
| Tree species richness (Sr) | 1 | 73 | 0.851 | 0.359 | 1.720 | 0.194 | 3.967 | *0.050* |
| Mycorrhizal type (Myc) | 2 | 73 | 1.772 | 0.177 | 1.585 | 0.212 | 2.338 | 0.104 |
| Sr x Myc | 2 | 73 | 1.146 | 0.324 | 0.422 | 0.658 | 0.187 | 0.830 |

Shown are the degrees of freedom (df), denominator degrees of freedom (ddf), *F*-values, and the statistical significance of the fixed effects (*p*-values). Significant effects ($p < 0.05$) are given in bold and marginally significant effects ($p < 0.1$) in italics.

Table S7 Summary of Tukey HSD analysis testing the difference in elemental contents among Mycorrhizal types (AM, EM, AM+EM; Myc) for (a) foliage, (b) soil, (c) soil microbial biomass.

| | | **C content** | | **N content** | | **P content** | |
|---|---|---|---|---|---|---|---|
| **Source of variation** | **df** | ***t*** | ***p*** | ***t*** | ***p*** | ***t*** | ***p*** |
| (a) Foliage | | | | | | | |
| AM vs. EM | 73 | -7.403 | **<.001** | -7.311 | **<.001** | 4.150 | **<.001** |
| AM vs. AM+EM | 73 | -2.725 | **0.022** | -2.302 | *0.062* | 1.632 | 0.238 |
| EM vs. AM+EM | 73 | 3.485 | **0.002** | 3.830 | **0.001** | -1.849 | 0.160 |
| (b) Soil | | | | | | | |
| AM vs. EM | 73 | -0.839 | 0.680 | -0.490 | 0.876 | 0.184 | 0.833 |
| AM vs. AM+EM | 73 | 0.335 | 0.940 | 0.411 | 0.911 | 0.305 | 0.583 |
| EM vs. AM+EM | 73 | 1.039 | 0.555 | 0.822 | 0.691 | 0.245 | 0.784 |
| (c) Soil microbial biomass | | | | | | | |
| AM vs. EM | 73 | -0.905 | 0.639 | -1.032 | 0.561 | -0.794 | 0.708 |
| AM vs. AM+EM | 73 | 0.241 | 0.968 | -0.608 | 0.816 | 0.922 | 0.628 |
| EM vs. AM+EM | 73 | 1.000 | 0.579 | 0.258 | 0.964 | 1.588 | 0.257 |

Shown are t-ratio and *p*-values for Tukey HSD analysis. Significant effects are given in bold and marginally significant effects in italics (*p*-values).

Table S8 Summary of Tukey HSD analysis testing the difference in elemental pools (carbon (C), nitrogen (N), phosphorus (P)) among Mycorrhizal types (AM, EM, AM+EM; Myc) for (a) foliage, (b) soil, (c) soil microbial biomass.

| | **C pool** | | **N pool** | | **P pool** | |
|---|---|---|---|---|---|---|
| **Source of variation** | ***t*** | ***p*** | ***t*** | ***p*** | ***t*** | ***p*** |
| (a) Foliage | | | | | | |
| AM vs. EM | -1.914 | 0.142 | -4.910 | **<.001** | 2.378 | *0.052* |
| AM vs. AM+EM | -1.237 | 0.435 | -2.071 | 0.103 | 0.474 | 0.884 |
| EM vs. AM+EM | 0.368 | 0.928 | 2.049 | 0.108 | -1.521 | 0.287 |
| (b) Soil | | | | | | |
| AM vs. EM | -1.537 | 0.28 | -1.279 | 0.411 | -0.717 | 0.754 |
| AM vs. AM+EM | 0.354 | 0.933 | 0.382 | 0.923 | 0.458 | 0.891 |
| EM vs. AM+EM | 1.643 | 0.234 | 1.455 | 0.318 | 1.060 | 0.542 |

| | | | | | | |
|---|---|---|---|---|---|---|
| (c) Soil microbial biomass | | | | | | |
| AM vs. EM | -1.363 | 0.366 | -1.555 | 0.271 | -0.915 | 0.633 |
| AM vs. AM+EM | 0.314 | 0.947 | -0.349 | 0.935 | 1.104 | 0.514 |
| EM vs. AM+EM | 1.457 | 0.318 | 0.956 | 0.607 | 1.872 | 0.154 |

Shown are t-ratio and *p*-values for Tukey HSD analysis. Significant effects are given in bold and marginally significant effects in italics (*p*-values).

Table S9 Summary of simple linear regression analyses between elemental content of (a) tree foliage (b), soil, and (c) soil microbial biomass and Tree species richness for AM, EM and AM+EM communities.

| | | **C content** | | **N content** | | **P content** | |
|---|---|---|---|---|---|---|---|
| **Source of variation** | **df** | ***F*** | ***p*** | ***F*** | ***p*** | ***F*** | ***p*** |
| (a) Foliage | | | | | | | |
| AM (n=30) | 1,28 | 0.74 | 0.397 | 7.205 | **0.012** | 7.75 | **0.01** |
| EM (n=30) | 1,28 | 0.018 | 0.895 | 4.504 | **0.043** | 0.271 | 0.607 |
| AM+EM (n=20) | 1,18 | 0.605 | 0.447 | 0.362 | 0.56 | 0.514 | 0.483 |
| (b) Soil | | | | | | | |
| AM (n=30) | 1,28 | 0.353 | 0.557 | 0.14 | 0.712 | 0.642 | 0.43 |
| EM (n=30) | 1,28 | 0.002 | 0.966 | 0.008 | 0.931 | 0.015 | 0.902 |
| AM+EM (n=20) | 1,18 | 0.004 | 0.952 | 0.077 | 0.785 | 0.057 | 0.813 |
| (c) Soil microbial biomass | | | | | | | |
| AM (n=30) | 1,28 | 0.274 | 0.604 | 3.247 | *0.082* | 6.979 | **0.013** |
| EM (n=30) | 1,28 | 3.978 | *0.056* | 2.036 | 0.165 | 3.62 | *0.067* |
| AM+EM (n=20) | 1,18 | <.001 | 0.993 | 0.012 | 0.913 | 0.067 | 0.798 |

Shown are the degrees of freedom (df), denominator degrees of freedom (ddf), *F*-values, and the statistical significance of the fixed effects (*p*-values). Significant effects ($p < 0.05$) are given in bold and marginally significant effects ($p < 0.1$) in italics.

Table S10 Summary of simple linear regression analyses between Tree species richness (Sr; one, two, four) of AM, EM and AM+EM communities of elemental pools (carbon (C), nitrogen (N), phosphorus (P)) within (a) tree foliage (b), soil, and (c) soil microbial biomass.

| | | C pool | | N pool | | P pool | |
|---|---|---|---|---|---|---|---|
| **Source of variation** | **df** | ***F*** | ***p*** | ***F*** | ***p*** | ***F*** | ***p*** |
| (a) Foliage | | | | | | | |
| AM (n=30) | 1,28 | 3.938 | *0.057* | 0.079 | 0.781 | 13.18 | **0.001** |
| EM (n=30) | 1,28 | 6.249 | **0.019** | 1.766 | 0.195 | 3.398 | *0.076* |
| AM+EM (n=20) | 1,18 | 3.851 | *0.065* | 1.292 | 0.271 | 1.817 | 0.194 |
| (b) Soil | | | | | | | |
| AM (n=30) | 1,28 | 0.177 | 0.677 | 0.262 | 0.613 | 0.037 | 0.849 |
| EM (n=30) | 1,28 | 0.005 | 0.942 | 0.001 | 0.974 | <.001 | 0.993 |
| AM+EM (n=20) | 1,18 | 0.097 | 0.758 | 0.126 | 0.727 | 0.069 | 0.795 |
| (c) Soil microbial biomass | | | | | | | |
| AM (n=30) | 1,28 | 0.01 | 0.923 | 0.813 | 0.375 | 3.819 | *0.061* |
| EM (n=30) | 1,28 | 2.905 | *0.099* | 1.36 | 0.253 | 2.5 | 0.125 |
| AM+EM (n=20) | 1,18 | 0.023 | 0.841 | 0.023 | 0.881 | 0.193 | 0.666 |

Shown are the degrees of freedom (df), denominator degrees of freedom (ddf), *F*-values, and the statistical significance of the fixed effects (*p*-values). Significant effects ($p < 0.05$) are given in bold and marginally significant effects ($p < 0.1$) in italics.

Table S11 Summary of RW1 (Random Walk model of order 1) model using Bayesian statistics testing the effects of Tree species richness (one, two, four; Sr), Mycorrhizal type (AM, EM, AM+EM; Myc), Season (Spring, Summer, Autumn, Winter), and Soil moisture (Moi) on (a) nitrate ($NO_3$-N), (b) ammonium ($NH_4$-N) and (c) phosphate ($PO_4$-P) availability.

| (a) **$NO_3$-N** | | | | |
|---|---|---|---|---|
| **parameter** | **mean** | **sd** | **2.5%** | **97.5%** |
| (Intercept) | **-1,52** | **0,57** | **-2,64** | **-0,42** |
| Tree species richness (Sr) | **-0,09** | **0,03** | **-0,15** | **-0,03** |
| Mycorrhizal type (Myc) AM + EM | -0,89 | 0,54 | -1,94 | 0,16 |
| Myc EM | -0,28 | 0,53 | -1,31 | 0,75 |
| Soil moisture (Moi) | 0,00 | 0,02 | -0,04 | 0,05 |
| Season Summer | -0,50 | 0,57 | -1,61 | 0,63 |
| Season Autumn | **-1,14** | **0,46** | **-2,04** | **-0,23** |
| Season Winter | **-1,80** | **0,53** | **-2,82** | **-0,75** |
| Myc AM + EM x Moi | 0,04 | 0,02 | -0,01 | 0,08 |
| Myc EM x Moi | 0,03 | 0,02 | -0,02 | 0,07 |
| Myc AM + EM x Season Summer | 0,09 | 0,22 | -0,34 | 0,53 |
| Myc EM x Season Summer | **-0,43** | **0,21** | **-0,84** | **0,00** |
| Myc AM + EM x Season Autumn | -0,05 | 0,18 | -0,41 | 0,30 |
| Myc EM x Season Autumn | **-0,54** | **0,18** | **-0,89** | **-0,19** |
| Myc AM + EM x Season Winter | 0,03 | 0,15 | -0,27 | 0,33 |
| Myc EM: Season Winter | -0,22 | 0,16 | -0,53 | 0,08 |
| Moi x Season Summer | 0,00 | 0,03 | -0,06 | 0,06 |
| Moi x Season Autumn | 0,03 | 0,02 | -0,01 | 0,07 |
| Moi x Season Winter | **0,05** | **0,02** | **0,01** | **0,09** |
| (b) **$NH_4$-N** | | | | |
| (Intercept) | **-2,88** | **0,42** | **-3,73** | **-2,07** |
| Tree species richness (Sr) | 0,00 | 0,01 | -0,02 | 0,02 |
| Mycorrhizal type (Myc) AM + EM | 0,17 | 0,34 | -0,50 | 0,84 |
| Myc EM | 0,23 | 0,30 | -0,36 | 0,83 |
| Soil moisture (Moi) | 0,01 | 0,02 | -0,02 | 0,04 |
| Season Summer | **-1,27** | **0,41** | **-2,07** | **-0,46** |
| Season Autumn | -0,62 | 0,32 | -1,26 | 0,01 |
| Season Winter | 0,36 | 0,34 | -0,30 | 1,03 |
| Myc AM + EM x Moi | -0,01 | 0,01 | -0,04 | 0,02 |
| Myc EM x Moi | -0,01 | 0,01 | -0,04 | 0,02 |
| Myc AM + EM x Season Summer | -0,04 | 0,13 | -0,30 | 0,21 |
| Myc EM x Season Summer | -0,02 | 0,12 | -0,25 | 0,20 |
| Myc AM + EM x Season Autumn | -0,05 | 0,12 | -0,28 | 0,19 |
| Myc EM x Season Autumn | 0,00 | 0,11 | -0,21 | 0,21 |
| Myc AM + EM x Season Winter | 0,01 | 0,10 | -0,19 | 0,21 |

| | | | | |
|---|---|---|---|---|
| Myc EM: Season Winter | 0,03 | 0,09 | -0,15 | 0,21 |
| Moi x Season Summer | **0,08** | **0,02** | **0,04** | **0,12** |
| Moi x Season Autumn | **0,02** | **0,01** | **0,00** | **0,05** |
| Moi x Season Winter | -0,02 | 0,01 | -0,04 | 0,01 |
| **(c) $PO_4$-P** | | | | |
| (Intercept) | **-8,15** | **0,88** | **-9,88** | **-6,45** |
| Tree species richness (Sr) | -0,03 | 0,04 | -0,10 | 0,05 |
| Mycorrhizal type (Myc) AM + EM | -0,79 | 0,81 | -2,39 | 0,81 |
| Myc EM | 0,21 | 0,70 | -1,16 | 1,58 |
| Soil moisture (Moi) | **0,15** | **0,03** | **0,09** | **0,22** |
| Season Summer | 0,91 | 0,96 | -0,98 | 2,79 |
| Season Autumn | -0,12 | 0,76 | -1,62 | 1,36 |
| Season Winter | 0,98 | 0,80 | -0,58 | 2,54 |
| Myc AM + EM x Moi | 0,03 | 0,03 | -0,04 | 0,10 |
| Myc EM x Moi | -0,01 | 0,03 | -0,07 | 0,05 |
| Myc AM + EM x Season Summer | 0,19 | 0,31 | -0,42 | 0,79 |
| Myc EM x Season Summer | -0,08 | 0,26 | -0,60 | 0,44 |
| Myc AM + EM x Season Autumn | 0,24 | 0,28 | -0,32 | 0,79 |
| Myc EM x Season Autumn | 0,00 | 0,24 | -0,48 | 0,47 |
| Myc AM + EM x Season Winter | 0,06 | 0,24 | -0,41 | 0,53 |
| Myc EM: Season Winter | 0,04 | 0,21 | -0,37 | 0,44 |
| Moi x Season Summer | -0,01 | 0,05 | -0,11 | 0,09 |
| Moi x Season Autumn | 0,03 | 0,03 | -0,03 | 0,10 |
| Moi x Season Winter | -0,04 | 0,03 | -0,10 | 0,02 |

Shown are the mean values (mean), standard deviation (sd), and the 2.5$^{th}$ and 97.5$^{th}$ percentiles of the parameter estimate values. Notably, parameter estimates are highlighted in bold if there is a 95% probability that they differ from zero.

Table S12 Summary of one-tailed t-test analyses (values higher than zero) for biodiversity effects (Net effects, Selection effects, and Complementarity effects) of tree foliage (a) C pool, (b) N pool, and (c) P pool for communities of AM, EM, or AM+EM trees with 2 tree species or 4 tree species.

| | | **Net effects** | | **Selection effects** | | **Complementarity effects** | |
|---|---|---|---|---|---|---|---|
| **Source of variation** | **df** | ***t*** | ***p*** | ***t*** | ***p*** | ***t*** | ***p*** |
| (a) C pool (n=10) | | | | | | | |
| AM | | | | | | | |
| 2-sp. | 9 | 3.623 | **0.006** | 4.313 | **0.002** | 2.582 | **0.030** |
| 4-sp. | 9 | 12.027 | **<.001** | 6.349 | **<.001** | 5.292 | **<.001** |
| EM | | | | | | | |
| 2-sp. | 9 | 1.395 | 0.197 | 2.910 | **0.020** | 0.244 | 0.813 |
| 4-sp. | 9 | 9.156 | **<.001** | 7.210 | **<.001** | 3.838 | **0.004** |
| AM+EM | | | | | | | |
| 2-sp. | 9 | 4.788 | **<.001** | 0.459 | 0.657 | 7.772 | **<.001** |
| 4-sp. | 9 | 11.751 | **<.001** | 8.603 | **<.001** | 9.917 | **<.001** |
| (b) N pool (n = 10) | | | | | | | |
| AM | | | | | | | |
| 2-sp. | 9 | 0.483 | 0.641 | 1.894 | *0.091* | 0.033 | 0.974 |
| 4-sp. | 9 | -0.600 | 0.563 | 1.894 | *0.091* | -1.095 | 0.302 |
| EM | | | | | | | |
| 2-sp. | 9 | 0.090 | 0.930 | 1.909 | *0.089* | -1.148 | 0.281 |
| 4-sp. | 9 | 2.667 | **0.026** | 5.036 | **0.001** | -0.906 | 0.389 |
| AM+EM | | | | | | | |
| 2-sp. | 9 | -0.227 | 0.826 | -3.598 | **0.006** | 0.942 | 0.371 |
| 4-sp. | 9 | 3.648 | **0.005** | 0.837 | 0.424 | 2.274 | **0.049** |
| (c) P pool (n = 10) | | | | | | | |
| AM | | | | | | | |
| 2-sp. | 9 | 0.583 | 0.574 | 0.405 | 0.695 | 0.233 | 0.821 |
| 4-sp. | 9 | 11.997 | **<.001** | 3.467 | **0.007** | 10.546 | **<.001** |

| | Net effects | | | Selection effects | | Complementarity effects | |
|---|---|---|---|---|---|---|---|
| Source of variation | df | *t* | *p* | *t* | *p* | *t* | *p* |
| EM | | | | | | | |
| 2-sp. | 9 | 1.488 | 0.171 | 4.409 | **0.002** | -0.933 | 0.375 |
| 4-sp. | 9 | 5.386 | **<.001** | 13.004 | **<.001** | 0.239 | 0.817 |
| AM+EM | | | | | | | |
| 2-sp. | 9 | 2.322 | **0.045** | 1.444 | 0.183 | 1.849 | *0.098* |
| 4-sp. | 9 | 5.972 | **<.001** | 4.492 | **0.002** | 2.890 | **0.017** |

Shown are degrees of freedom (df), *t*- and *p*-values, and plot number (n). Significant effects are given in bold and marginally significant effects in italics (*p*-values).

Table S13 Summary of Tukey HSD analysis testing the difference in biodiversity effects (Net effects, Selection effects, and Complementarity effects) among Mycorrhizal types (AM, EM, AM+EM; Myc) for (a) carbon (C), (b) nitrogen (N), and (c) phosphorous (P) pools.

| | Net effects | | Selection effects | | Complementarity effects | |
|---|---|---|---|---|---|---|
| Source of variation | *t* | *p* | *t* | *p* | *t* | *p* |
| (a) C pool | | | | | | |
| AM vs. EM | -0.569 | 0.837 | -2.234 | *0.075* | 0.940 | 0.618 |
| AM vs. AM+EM | -1.080 | 0.530 | 0.634 | 0.802 | -1.877 | 0.155 |
| EM vs. AM+EM | -0.511 | 0.866 | 2.868 | **0.016** | -2.817 | **0.018** |
| (b) N pool | | | | | | |
| AM vs. EM | -1.295 | 0.404 | -2.330 | *0.060* | 0.653 | 0.792 |
| AM vs. AM+EM | -1.058 | 0.544 | 2.286 | *0.066* | -1.840 | 0.166 |
| EM vs. AM+EM | 0.237 | 0.970 | 4.616 | **<.001** | -2.493 | **0.041** |
| (c) P pools | | | | | | |
| AM vs. EM | 0.290 | 0.955 | -4.675 | **<.001** | 3.036 | **0.010** |
| AM vs. AM+EM | -0.583 | 0.830 | -1.921 | 0.143 | 0.040 | 0.999 |
| EM vs. AM+EM | 0.873 | 0.660 | 2.754 | **0.022** | -2.995 | **0.011** |

Shown are *t*- and *p*-values for Tukey HSD analysis. Significant effects are given in bold and marginally significant effects in italics (*p*-values).

Table S14 Summary of simple linear regression analyses between biodiversity effects (Net effects, Selection effects, and Complementarity effects) of tree foliage and Tree species richness of AM, EM, and AM+EM communities.

| | | Net effects | | Selection effects | | Complementarity effects | |
|---|---|---|---|---|---|---|---|
| **Source of variation** | **df** | ***F*** | ***p*** | ***F*** | ***p*** | ***F*** | ***p*** |
| (a) C pool | | | | | | | |
| AM (n = 20) | 1, 18 | 1.401 | 0.252 | 0.858 | 0.367 | 0.615 | 0.443 |
| EM (n = 20) | 1, 18 | 4.977 | **0.039** | 3.826 | *0.066* | 2.943 | 0.103 |
| AM+EM (n = 20) | 1, 18 | 7.046 | **0.016** | 22 | **<.001** | 0.967 | 0.339 |
| (b) N pool | | | | | | | |
| AM (n = 20) | 1, 18 | 0.540 | 0.472 | 0.021 | 0.887 | 0.472 | 0.501 |
| EM (n = 20) | 1, 18 | 2.138 | 0.161 | 1.915 | 0.183 | 0.172 | 0.683 |
| AM+EM (n = 20) | 1, 18 | 3.968 | *0.062* | 6.899 | **0.017** | 0.113 | 0.740 |
| (c) P pool | | | | | | | |
| AM (n = 20) | 1, 18 | 11.57 | **0.003** | 1.896 | 0.185 | 12.56 | **0.002** |
| EM (n = 20) | 1, 18 | 2.737 | 0.115 | 5.994 | **0.025** | 0.752 | 0.397 |
| AM+EM (n = 20) | 1, 18 | 3.379 | *0.083* | 5.896 | **0.026** | 0.352 | 0.560 |

Shown are degrees of freedom (df; numerator degrees of freedom), *F*- and *p*-values, and plot number (n). Significant effects are given in bold and marginally significant effects in italics (*p*-values).

Table S15 Percentage changes showing the strength of effects of Tree species richness (one, two, four; Sr) on elemental contents in (a) foliage, (b) soil, and (c) microbial biomass.

| | **Sr 1 vs. Sr 2** | **Sr 1 vs. Sr 4** | **Sr 2 vs. Sr 4** |
|---|---|---|---|
| (a) Foliage elemental contents | | | |
| C content | -0.001 | -0.005 | -0.005 |
| N content | -0.092 | -0.132 | -0.045 |
| P content | 0.014 | 0.143 | 0.127 |
| (b) Soil | | | |
| C content | 0.023 | 0.009 | -0.013 |
| N content | 0.008 | 0.003 | -0.005 |
| P content | 0.013 | 0.015 | 0.002 |
| (c) Soil microbial biomass | | | |
| C content | 0.004 | 0.012 | 0.008 |
| N content | 0.017 | 0.030 | 0.013 |
| P content | -0.112 | 0.182 | 0.331 |

Table S16 Percentage changes showing the strength of effects of Tree species richness (one, two, four; Sr) on (a) elemental pools in (a) foliage, (b) soil, and (c) microbial biomass.

| | **Sr 1 vs. Sr 2** | **Sr 1 vs. Sr 4** | **Sr 2 vs. Sr 4** |
|---|---|---|---|
| (a) Foliage | | | |
| C pool | 0.121 | 0.246 | 0.111 |
| N pool | 0.013 | 0.086 | 0.072 |
| P pool | 0.141 | 0.407 | 0.233 |
| (b) Soil | | | |
| C pool | 0.009 | -0.027 | -0.036 |
| N pool | -0.007 | -0.034 | -0.027 |
| P pool | -0.005 | -0.023 | -0.018 |
| (c) Soil microbial biomass | | | |
| C pool | 0.006 | 0.014 | 0.008 |
| N pool | 0.019 | 0.028 | 0.009 |
| P pool | -0.061 | 0.358 | 0.444 |

Table S17 Percentage changes showing the strength of effects of Mycorrhizal type (AM, EM, AM+EM, Myc) on elemental contents in (a) foliage, (b) soil, and (c) microbial biomass.

| | **AM vs. EM** | **AM vs. AM + EM** | **EM vs. AM + EM** |
|---|---|---|---|
| (a) Foliage | | | |
| C content | -0.051 | -0.021 | 0.030 |
| N content | -0.244 | -0.082 | 0.162 |
| P content | 0.231 | 0.072 | -0.158 |
| (b) Soil | | | |
| C content | -0.018 | 0.008 | 0.026 |
| N content | -0.009 | 0.011 | 0.019 |
| P content | 0.012 | 0.015 | 0.003 |
| (c) Soil microbial biomass | | | |
| C content | -0.006 | 0.002 | 0.007 |
| N content | -0.012 | -0.011 | 0.001 |
| P content | -0.073 | 0.152 | 0.243 |

Table S18 Percentage changes showing the strength of effects of Mycorrhizal type (AM, EM, AM+EM, Myc) on (a) elemental pools in (a) foliage, (b) soil, and (c) microbial biomass.

| | **AM vs. EM** | **AM vs. AM + EM** | **EM vs. AM + EM** |
|---|---|---|---|
| (a) Foliage | | | |
| C pool | -0.088 | -0.114 | -0.026 |
| N pool | -0.278 | -0.178 | 0.100 |
| P pool | 0.184 | -0.030 | -0.214 |
| (b) Soil | | | |
| C pool | -0.062 | 0.027 | 0.095 |
| N pool | -0.053 | 0.030 | 0.068 |
| P pool | -0.034 | 0.035 | 0.071 |
| (c) Soil microbial biomass | | | |
| C pool | -0.017 | 0.007 | 0.024 |
| N pool | -0.028 | -0.008 | 0.020 |
| P pool | -0.145 | 0.218 | 0.425 |
